# Identification of novel thiazole derivatives as flaviviral protease inhibitors against DENV and JEV

**DOI:** 10.1101/2024.11.04.621919

**Authors:** Sheikh Murtuja, Sayan Das, Indrani Das Jana, Deepak Shilkar, Gourav Rakshit, Biswatrish Sarkar, Barij Nayan Sinha, Rikeshwar Prasad Dewangan, Arindam Mondal, Venkatesan Jayaprakash

**Affiliations:** Department of Pharmaceutical Sciences & Technology, Birla Institute of Technology, Mesra, Ranchi 835215, Jharkhand, India; Department of Bioscience and Biotechnology, Indian Institute of Technology, Kharagpur 721302, West Bengal, India; Department of Pharmaceutical Chemistry, School of Pharmaceutical Education and Research, Jamia Hamdard, New Delhi110062

**Author notes:** Corresponding Authors: Arindam Mondal (AM), Venkatesan Jayaprakash (VJ). The authors have contributed equally to this work (Sheikh Murtuja, Sayan Das).

**Keywords:** Antivirals, NS2B-NS3 Protease inhibitors, Thiazoles, Dengue virus, Japanese encephalitis virus

## Abstract

Flaviviruses are the causative agents of viral hemorrhagic fever (VHF) globally and have demonstrated the capacity to result in fatal outcomes if not managed effectively. Among different types of flaviviruses, dengue (DENV) and Japanese encephalitis (JEV) are the most common in tropical and subtropical countries. Given the scarcity of effective vaccines against flaviviral infections, small molecule agents that target crucial viral proteins emerge as the sole feasible option. This study outlines the synthesis of novel thiazole compounds as flavivirus NS2B-NS3 protease inhibitor and characterization of their antiviral activity against DENV and JEV. We synthesized a heterocyclic template derived from a substrate-based retrotripeptide dengue protease inhibitor, leading to forty-eight thiazole derivatives. Two compounds demonstrated significant inhibition of dengue virus protease activity *in vitro*. Comprehensive characterization of these two compounds was conducted through biochemical assay which revealed an uncompetitive mode of inhibition. Subsequent cell-based assays using Dengue and Japanese encephalitis viruses as representatives Flaviviruses imparted the potential of these compounds to block viral RNA synthesis, and viral replication. Finally time-course experiments unveiled that the two compounds impeded the accumulation of viral genomic RNA primarily at later stages of infection, aligning with their capacity to hinder NS2B-NS3 protease activity, polyprotein processing and viral genomic RNA replication. Together our work presents the development and validation of flavivirus protease inhibitors with therapeutic potential against Dengue and Japanese encephalitis virus.

## Introduction

Flaviviruses belong to the Flaviviridae family and comprise nearly 70 pathogenic viruses, including dengue virus (DENV), Japanese encephalitis virus (JEV), West Nile virus (WNV), Yellow fever virus (YFV), and Zika virus (ZIKV).(1, 2) These viruses are primarily arthropod-borne (arboviruses), transmitted through mosquitoes or ticks, and poses significant threat to global human health and economy(3, 4). Members of this genus are of great concern, as they play a significant role in the emergence of severe infectious diseases. The rising prevalence of DENV in tropical and subtropical regions, the occurrence of WNV in North America, and the spread of JEV across a substantial part of Asia and Oceania remain pressing issues(5). DENV is the most common arbovirus and is endemic to approximately 110 countries, with an estimated 400 million infections and 20,000 deaths annually(6). DENV is an enveloped positive-sense RNA virus that causes dengue fever (DF), dengue hemorrhagic fever (DHF), and dengue shock syndrome (DSS) in humans(6, 7). To date, there has been no officially sanctioned antiviral therapy, and the vaccine Dengvaxia, the only one granted regulatory approval, has not demonstrated any encouraging outcomes in the context of mitigating dengue epidemics(8). Rather, this vaccine elicits significant worries about antibody-dependent enhancement of illness (ADE)(9–11). Therefore, developing novel therapeutic approaches for DENV infections or creating broad-spectrum inhibitors for closely related flaviviruses is of utmost urgency.(12)

DENV infection starts with receptor-mediated endocytosis of the virus particle, followed by the deposition of viral genetic material within the cytoplasm(6, 7). The genomic material of the virus is a 10.7-kilobase, capped, positive-sense RNA that is translated into a single polyprotein upon its release on the endoplasmic reticulum (ER) membrane. Both host and viral-encoded proteases subsequently process this polyprotein into three structural proteins, namely the capsid (C), membrane (M), and envelope (E) proteins, as well as seven non-structural proteins, namely NS1, NS2A, NS2B, NS3, NS4A, NS4B, and NS5(13–15). These newly synthesized viral proteins stimulate the transition of the viral genome to be used as a substrate for translation to a template for genomic RNA replication(16). Non-structural proteins, together with multiple host factors, constitute viral replication complexes that execute viral genomic RNA replication within infection- induced double membrane vesicle packets (VP) generated on the ER membranes(15). The newly synthesized genomic RNA was packaged with different structural proteins along the host-derived membrane to generate premature and mature virion particles(15). The non-structural proteins NS2B and NS3 together reconstitute the viral serine protease, which is instrumental for polyprotein processing in the early phase of infection and virion maturation at later stages, thus acting as a critical modulator of the progression of infection(17). In the NS2B-NS3 complex, the NS3 protein constitutes the catalytic core, whereas NS2B acts as a cofactor important for stabilization and substrate recognition(18). The N-terminal domain of NS3 contains a highly conserved protease- active core consisting of His51, Asp75, and Serl35 triad motifs that are critical for enzymatic activity(19, 20). The NS2B-NS3 protease remains an attractive target for antiviral drug development due to its role in flaviviral replication(21).

Viral proteases are well-accepted as effective targets for the discovery and development of antivirals(22). The clinical effectiveness of viral protease inhibitors has been established by incorporating these inhibitors as major components in antiretroviral therapy(23, 24). Although substrate-based peptidomimetics dominate protease inhibitor design, the low stability and bioavailability of peptide-based drugs have prompted the need for non-peptidic, small-molecule-based protease inhibitors(25, 26). Peptidic and non-peptidic inhibitors of DENV NS2B-NS3 reported were reviewed from time to time by our group (27–29) and other scientific groups(30–33).

Here we present the design, synthesis, and validation of two novel DENV protease inhibitors by transforming a retrotripeptide, previously reported by Weigel et al. 2015. Flavivirus proteases prefer substrates containing dibasic amino acids (Lys or Arg) at the P1 and P2 positions on the non-prime side(34). Weigel *et al.* addressed this requirement by retaining lysine at the P1 position and replacing arginine at the P2 position with its mimic, *p*-acetamidinyl phenyl ring. In our design process, we evaluated the possibility of integrating an oxazole-based heterocyclic template on the N-terminal side of this retro-tripeptide (see Figure 1). The design incorporates a 2-phenyl oxazole with substitutions at the 4^th^ and 5^th^ positions, strategically placed in the S1 and S2 sub- pockets of the enzyme, respectively. We replaced the oxazole with its ring counterpart, thiazole, to produce a thiazole derivative known for its extensively documented antiviral properties(35, 36). Moreover, substitution at the second position of these thiazole molecules might occupy either the S1 or S2 sub-pockets due to rotational flexibility around the single bond linking the phenyl ring to the thiazole ring (Fig. 1).

**Figure 1.**
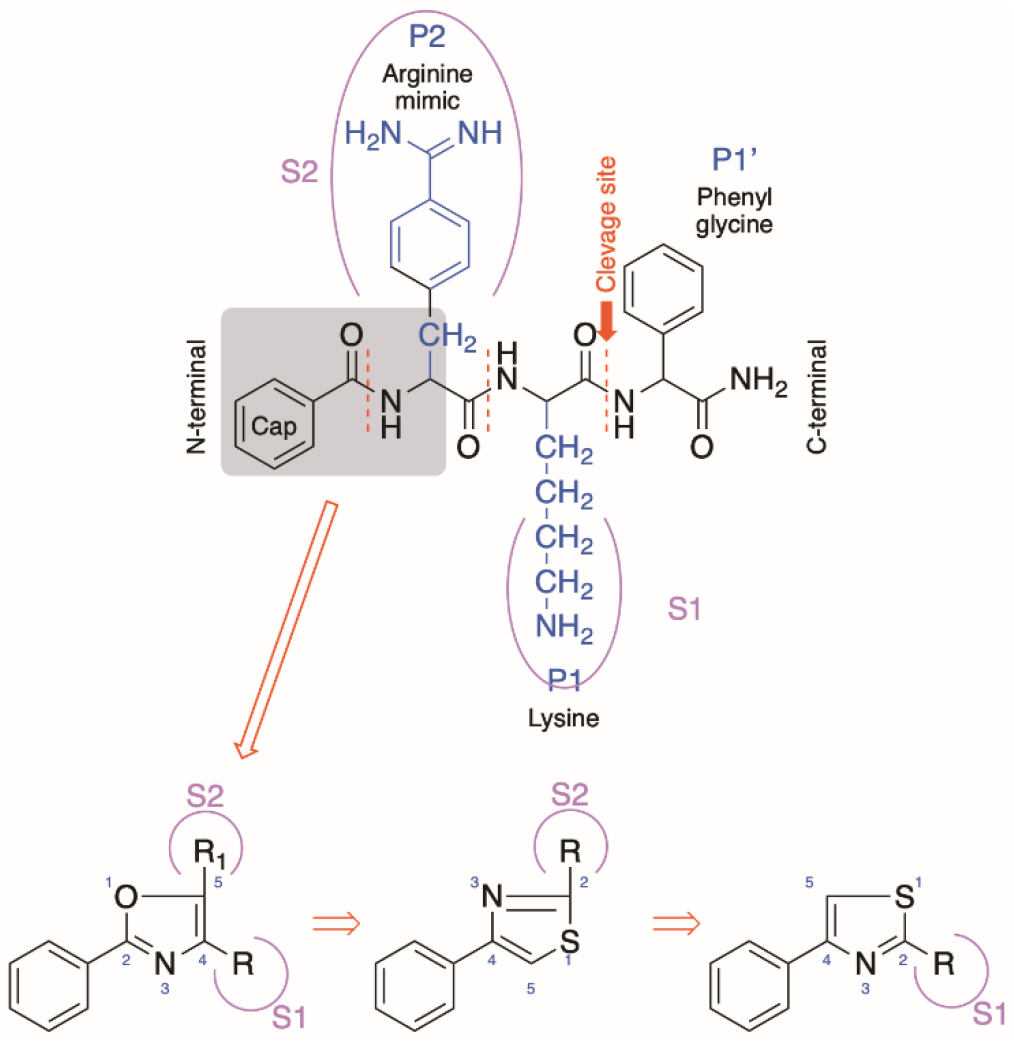
The strategy of designing small molecule inhibitors against Dengue protease

We synthesized forty-eight thiazole derivatives and conducted *in vitro* screening for their activity against DENV NS2B-NS3 protease. Two lead compounds featuring –OH and – OCH_3_ substitutions and displaying IC_50_ values below ∼25 µM were selected for further investigation regarding their antiviral efficacy against live DENV2 and JEV infections. Time-course experiments conclusively demonstrated that the compounds inhibited viral RNA replication solely at later stages of infection, proving their ability to inhibit protease activity, polyprotein processing, and the production of non-structural proteins.

Furthermore, we evaluated the anti-JEV effects of these compounds in SHSY-5Y cells to strengthen its potential as anti-Flaviviral drug.

## Results and Discussion Synthesis of thiazole derivatives

A set of forty-eight thiazole derivatives (**3a**–**3av**) was synthesized through the reactions illustrated in Scheme 1. Suitably substituted benzaldehydes and acetophenones (**1a– 1u**) were subjected to reaction with thiosemicarbazide in the presence of a catalytic quantity of glacial acetic acid in methanol. The reaction was conducted at room temperature with continuous stirring for about 4–6 h. The resulting thiosemicarbazones (**2a–2u**) were subsequently treated with appropriately substituted phenacyl bromide in methanol. The reaction was performed at room temperature under constant stirring for approximately 0.5-2 h. The resultant thiazole derivatives (**3a–3av**) were purified through recrystallization from hot methanol. All final compounds were characterized by ^1^H-NMR, ^13^C-NMR, and ESI-MS spectroscopy. The protons of the methyl group (-CH_3_) at positions R (**3a**, **3b**, **3j**, **3s**, **3t**) and R_1_ (**3s**–**3av**) appeared as singlets (s) with chemical shifts in the range of δ2.18–2.65 ppm. Additionally, a singlet (s) was observed in the range of δ3.74–3.95 ppm for the protons of the methoxy (-OCH_3_) group at R (**3c–3e, 3l– 3n, 3w–3ab, 3ah–3ak**) and R_2_ (**3aq, 3au**). The aromatic protons, including those in the thiazole ring (Ar-H), displayed multiplets in the range of δ6.20–9.91 ppm. Moreover, the vinylic proton at the R_1_ position (**3a–3r**) was detected in the aromatic region, coalescing with the aromatic proton peaks. A chemical shift ranging from δ9.03–9.87 ppm was observed for the proton of the -OH groups at R (**3u, 3v, 3af, 3ag, 3ao–3av**), appearing as a singlet. Additionally, for all compounds (**3a–3av**), the side chain -NH proton prominently appeared as a singlet within the range of δ10.96–12.56 ppm. ^13^C-NMR of compounds displayed peaks in the range of δ14.4–55.6 ppm and δ99.2–171.0 ppm for aliphatic and aromatic carbons, respectively. All the compounds exhibited either [M]^+^ or [M+1]^+^ as the base peak in ESI-MS. For halogenated compounds, [M+2]^+^ peaks were also detected. The spectra of all compounds are provided in the *Supplementary material*.

**Scheme 1.**
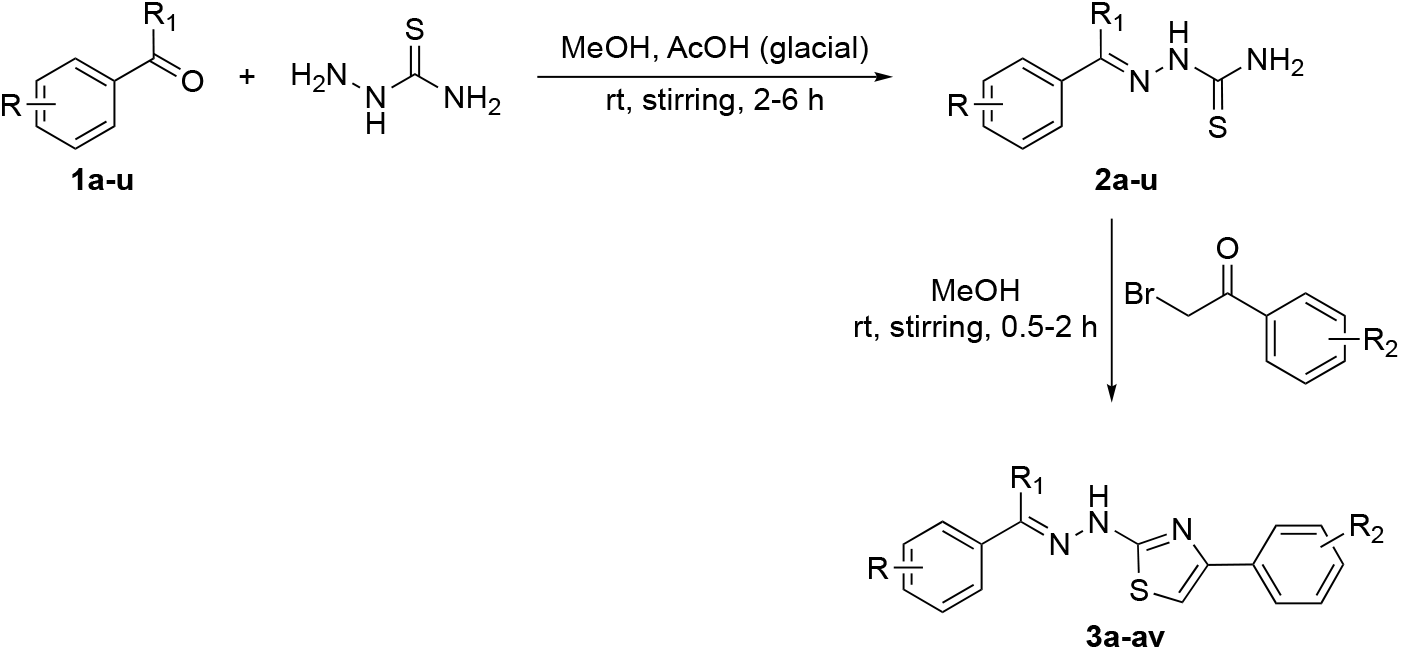
Synthesis of 2-[(2E)-2-benzylidene/-(1-phenylethylidene)-hydrazinyl]-4- phenyl-1,3-thiazole

### *In vitro* screening for anti-flaviviral protease activity

The synthesized compounds were screened to assess their ability to inhibit the flaviviral protease NS2B-NS3 activity. NS2B-NS3 displays trypsin-like serine protease activity and plays a role in cleaving the flavivirus polyprotein at the NS2A-NS2B, NS2B-NS3, NS3-NS4A, and NS4B-NS5 junctions(37). All flavivirus (DENV, JEV, YFV, WNV, and ZIKV) NS2B-NS3 proteases were structurally conserved despite their low primary sequence homology (Fig. 2A). Additionally, the catalytic core contains entirely conserved residues, including His51, Asp75, and Ser135 (according to the DENV NS3 protease residue position), arranged in a catalytic triad with identical spatial organization (Fig. 2B). This configuration represents an optimal target for the development of broad-spectrum NS2B-NS3 protease inhibitors.

**Figure 2.**
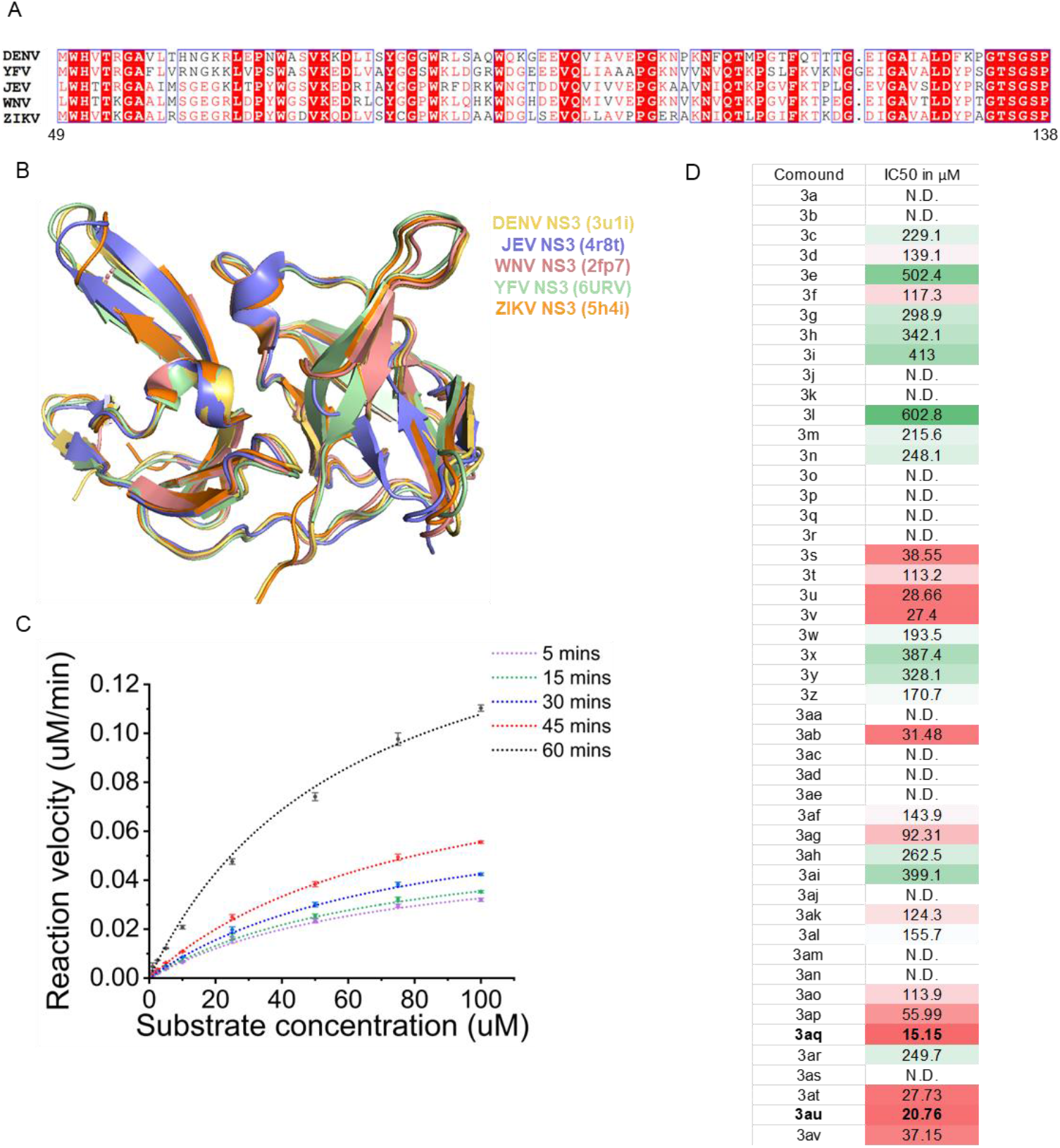
Conservation of the catalytic core region of NS3 protease and In vitro screening for anti-NS2B-NS3 protease activity (A) Multiple sequence alignment of NS3 catalytic domain of different flaviviruses, DENV (PDB ID: 3U1I), YFV (6URV), JEV (4R8T), WNV NS3 (2FP7), & ZIKV NS3 (5H4I). (B) Superimposed images of the NS3 protease of different flaviviruses showing its structural conservation. (C) NS2B-NS3 enzyme kinetics at different time points using increasing concentrations of fluorogenic substrate. (D) *In vitro* DENV2 NS2B-NS3 protease inhibition assay of forty-eight synthesised thiazole compounds at different concentrations. IC_50_ was determined by fitting nonlinear equation in Origin2023b software. Heatmap showing the IC_50_ of the compounds.

To assess the anti-DENV NS2B-NS3 activity of the compounds, we expressed the DENV2 NS2B-NS3 protease in BL21DE3 cells and subsequently purified it using Ni- NTA affinity chromatography, as outlined in the Methods section(38, 39). The protease activity assay was performed using different concentrations (1, 2.5, 5, 10, 25, 50, 75, and 100 μM) of a fluorogenic substrate (Abs-Nle-Lys-Arg-Arg-Ser-3-(NO_2_)Tyr) at various time points (5, 15, 30, 45, and 60 min), as described previously(38, 40). In our assay, the purified NS2B-NS3 protease showed a notable dose-dependent increase in reaction velocity after 1 h, with a less pronounced enhancement observed for shorter incubation times (Fig. 2C). Therefore, all subsequent assays were conducted with a 1- hour incubation time. The optimized reaction was then replicated with different concentrations of compounds (**3a**–**3av)** to deduce the half maximal inhibitory concentration (IC_50_). Of the forty-eight thiazole compounds tested, five compounds (3u, 3v, 3aq, 3au, and 3at) showed significant inhibition against DENV2 NS2B-NS3 protease with an IC_50_ value of less than 30 μM. Among them, compounds **3au** and **3aq** showed the most promising results, with an IC_50_ value of 20.76 μM and 15.15 μM respectively, compared to the dimethyl sulfoxide (DMSO) control (Fig. 2D).

## Structure-activity relationships (SAR)

To enable an SAR discussion, the data obtained from the *in vitro* screening are presented in Table 1–3. The activity data for benzaldehyde-based thiazole derivatives (**3a–3r**) are presented in Table 1. For the 3-nitro derivatives (**3a–3i**), only **3d** and **3f,** featuring a 2,5-diOCH_3_ and 2-F substitutions, showed IC_50_ value of 139.1 and 117.3 µM, respectively. Among the 4-Cl derivatives (**3j–3r**), none of the compound showed IC_50_ value less than 150 µM. It appears that electron-donating substitution (–OCH_3_) at the *ortho* (*2*) and *meta* (*5*) positions (**3d**) and -F substitution at the *ortho* (*2*) position (**3F**) in the 3-NO_2_ derivatives are more favourable than that in the 4-Cl derivative. However, other substitutions in both 3-NO_2_ and 4-Cl derivatives have shown IC_50_ value above 200 µM.

**Table 1.** DENV2 NS2B-NS3 protease inhibitory activity for Compounds 3a-3r

| Code | R | R<sub>1</sub> | R<sub>2</sub> | IC<sub>50</sub> (μM) |
| --- | --- | --- | --- | --- |
| 3a | 4-CH<sub>3</sub> | H | 3-NO<sub>2</sub> | ND |
| 3b | 3,5- (CH<sub>3</sub>)<sub>2</sub> | H | 3-NO<sub>2</sub> | ND |
| 3c | 2-OCH<sub>3</sub> | H | 3-NO<sub>2</sub> | 229.1 |
| 3d | 2,5-(OCH<sub>3</sub>)<sub>2</sub> | H | 3-NO<sub>2</sub> | 139.1 |
| 3e | 3,4-(OCH<sub>3</sub>)<sub>2</sub> | H | 3-NO<sub>2</sub> | 502.4 |
| 3f | 2-F | H | 3-NO<sub>2</sub> | 117.3 |
| 3g | 3-Cl | H | 3-NO<sub>2</sub> | 298.9 |
| 3h | 2-Br | H | 3-NO<sub>2</sub> | 342.1 |
| 3i | 3-Br | H | 3-NO<sub>2</sub> | 413 |
| 3j | 4-CH<sub>3</sub> | H | 4-Cl | 602.8 |
| 3k | 3,5- (CH<sub>3</sub>)<sub>2</sub> | H | 4-Cl | ND |
| 3l | 2-OCH<sub>3</sub> | H | 4-Cl | ND |
| 3m | 2,5-(OCH<sub>3</sub>)<sub>2</sub> | H | 4-Cl | 215.6 |
| 3n | 3,4-(OCH<sub>3</sub>)<sub>2</sub> | H | 4-Cl | 248.1 |
| 3o | 2-F | H | 4-Cl | ND |
| 3p | 3-Cl | H | 4-Cl | ND |
| 3q | 2-Br | H | 4-Cl | ND |
| 3r | 3-Br | H | 4-Cl | ND |

Among the thiazole derivatives of acetophenone origin (**3s-3an,** Table 2), we compared seven thiazole analogs of benzaldehyde origin (**3a**, **3c**, **3d**, **3e**, **3j**, **3m**, and **3n**) with those of acetophenone origin (**3t**, **3w**, **3aa**, **3ab**, **3ae**, **3ak**, and **3al**). The substitution of - H with -CH_3_ at the R_1_ position, we observed significant improvement in potency in case of both 3-NO2 (**3a** and **3e**) and 4-Cl (**3ak** and **3al**) derivatives. A ∼16-fold improvement was observed with **3ab** in comparison with **3e**. It is interesting to note that **3d** showing activity at 139.1 µM, completely lost its activity due to substitution of -H with -CH_3_ at R_1_ position (**3aa**). Whereas, **3e** showing activity ∼500 µM has shown dramatic increase in potency due to similar modification (**3ab**, 31.38 µM). This clearly shows that steric factors play a major role in accommodating the compound within the pocket with different binding conformation. This fact required to be checked with simulation studies. Compounds **3ak** and **3al** (4-Cl analogues of **3aa** and **3ab**, respectively) where found to be ∼1-fold better than their benzaldehyde counterparts, **3m** and **3n**. We expect no change in binding conformation with these compounds except improved hydrophobic interaction for compound **3ak** and **3al** due to -CH_3_ substitution at R_1_.

**Table 2.** DENV protease inhibitory activity for Compounds 3s-3an.

| Code | R | R<sub>1</sub> | R<sub>2</sub> | IC<sub>50</sub> (μM) |
| --- | --- | --- | --- | --- |
| 3s | 2-CH<sub>3</sub> | CH<sub>3</sub> | 3-NO<sub>2</sub> | 38.55 |
| 3t | 4-CH<sub>3</sub> | CH<sub>3</sub> | 3-NO<sub>2</sub> | 113.2 |
| 3u | 4-OH | CH<sub>3</sub> | 3-NO<sub>2</sub> | 28.66 |
| 3v | 3,5-(OH)<sub>2</sub> | CH<sub>3</sub> | 3-NO<sub>2</sub> | 27.4 |
| 3w | 2-OCH<sub>3</sub> | CH<sub>3</sub> | 3-NO<sub>2</sub> | 193.5 |
| 3x | 3-OCH<sub>3</sub> | CH<sub>3</sub> | 3-NO<sub>2</sub> | 387.4 |
| 3y | 4-OCH<sub>3</sub> | CH<sub>3</sub> | 3-NO<sub>2</sub> | 328.1 |
| 3z | 2,4-(OCH<sub>3</sub>)<sub>2</sub> | CH<sub>3</sub> | 3-NO<sub>2</sub> | 170.7 |
| 3aa | 2,5-(OCH<sub>3</sub>)<sub>2</sub> | CH<sub>3</sub> | 3-NO<sub>2</sub> | ND |
| 3ab | 3,4-(OCH<sub>3</sub>)<sub>2</sub> | CH<sub>3</sub> | 3-NO<sub>2</sub> | 31.38 |
| 3ac | 4-Cl | CH<sub>3</sub> | 3-NO<sub>2</sub> | ND |
| 3ad | 4-F | CH<sub>3</sub> | 3-NO<sub>2</sub> | ND |
| 3ae | 4-CH<sub>3</sub> | CH<sub>3</sub> | 4-Cl | ND |
| 3af | 4-OH | CH<sub>3</sub> | 4-Cl | 143.9 |
| 3ag | 3,5-(OH)<sub>2</sub> | CH<sub>3</sub> | 4-Cl | 92.31 |
| 3ah | 3-OCH<sub>3</sub> | CH<sub>3</sub> | 4-Cl | 262.5 |
| 3ai | 4-OCH<sub>3</sub> | CH<sub>3</sub> | 4-Cl | 399.1 |
| 3aj | 2,4-(OCH<sub>3</sub>)<sub>2</sub> | CH<sub>3</sub> | 4-Cl | ND |
| 3ak | 2,5-(OCH<sub>3</sub>)<sub>2</sub> | CH<sub>3</sub> | 4-Cl | 124.3 |
| 3al | 3,4-(OCH<sub>3</sub>)<sub>2</sub> | CH<sub>3</sub> | 4-Cl | 155.7 |
| 3m | 4-Cl | CH<sub>3</sub> | 4-Cl | ND |
| 3an | 4-F | CH<sub>3</sub> | 4-Cl | ND |

It is also observed that monomethoxy substituted derivatives (**3w**-**3y**) exhibited activity between ∼200 -∼400 µM concentration. Whereas their dimethoxy analogues (**3z**-**3ab**) displayed distance dependent activity. When they are *ortho* to each other displayed the best activity (**3ab**, 31.38 µM), *meta* to each other displayed moderate activity (**3z**, 170.7 µM), and *para* to each other displayed no activity (**3aa**, ND). We expect favorable hydrogen bonding interaction(s) for compound **3ab** as reason for its better activity profile. Accordingly, compounds **3u** (28.66 µM) and **3v** (27.40 µM) displayed potency equivalent to compound **3ab** (31.38 µM) as both were having -OH functional group capable of establishing H-bonding interaction with target protein. A similar pattern was observed with 4-Cl counterparts (**3af**-**3al**), but are less potent.

Considering the activity of **3u** and **3v**, we retained the substitution at R and R_1_ and went on to vary the substitution at R_2_ (Table 3). Substitution at para position with methoxy group provided the potent compounds **3aq** and **3au** of the series with IC_50_ value of 15.15 and 20.76 µM, respectively. Methyl and Fluro are Grimm’s hydride isosteres with exactly opposite electronic characteristics. Activity profile of these derivatives displayed no significant change in potency with respect to 3,5-dihydroxy analogues (**3at** and **3av**) but exhibited >4-fold difference in case of 4-hydroxy analogues (**3ap** and **3ar**). This once again shows that rather than electronic parameter, steric factor plays a role and accordingly we expect a different binding orientation for 3aq and 3au withing the binding pocket of DENV NS2B-NS3.

**Table 3.** DENV protease inhibitory activity for Compounds 3ao-3av

| Code | R | R<sub>1</sub> | R<sub>2</sub> | IC<sub>50</sub> (μM) |
| --- | --- | --- | --- | --- |
| 3ao | 4-OH | CH<sub>3</sub> | H | 113.9 |
| 3ap | 4-OH | CH<sub>3</sub> | 4-CH<sub>3</sub> | 55.99 |
| **3aq** | 4-OH | CH<sub>3</sub> | 4-OCH<sub>3</sub> | 15.15 |
| 3ar | 4-OH | CH<sub>3</sub> | 4-F | 249.7 |
| 3as | 3,5-(OH)<sub>2</sub> | CH<sub>3</sub> | H | ND |
| 3at | 3,5-(OH)<sub>2</sub> | CH<sub>3</sub> | 4-CH<sub>3</sub> | 27.73 |
| **3au** | 3,5-(OH)<sub>2</sub> | CH<sub>3</sub> | 4-OCH<sub>3</sub> | 20.76 |
| 3av | 3,5-(OH)<sub>2</sub> | CH<sub>3</sub> | 4-F | 37.15 |

## Interaction profile of the antiviral compound with the catalytic core

To elucidate the crucial interactions underlying DENV protease inhibitory activity, binding mode analysis was conducted for the top two compounds (**3aq** and **3au**).

Structure of active closed NS2B-NS3 complex(41) of DENV3 (PDB: 3U1I)(42) was used for the interaction study (Fig . 3A & 3B). In order to mimic the binding pocket of DENV2 the following residues were mutated: V36I, T115L, and K157R. Both the compounds, **3aq** and **3au** accommodated within the pocket in a similar fashion and displayed interactions that are common for both (Fig. 3A). Hydroxy functional group (4-OH & 3,5- (OH)_2_) established three H-bonding interaction with sidechain carboxylic acid group of ASP129 (S3), backbone carbonyl oxygen of PHE130, and sidechain phenolic hydroxyl group of TYR150 (S1). Anchoring of hydroxyl functional group of **3aq** and **3au** due to these H-bonds facilitate hydrophobic interaction of phenyl ring and R1 methyl group with S1’ (HIS51, SER135, GLY151) and S3 (ASP129) subpocket residues. Both S1, S1’ and S3 were not occupied rather the peripheral region (entrance) shows interaction with **3aq** and **3au**, interaction like substrate peptide backbone. An additional H-bonding interaction was observed between sidechain phenolic hydroxyl group with TYR161 with sidechain imino nitrogen of **3aq.** In case of **3au**, presence of dihydroxy functional group slightly pushed the compound that facilitated establishment of two additional H-bonding interaction for **3au**. One between phenolic hydroxyl group of TYR161 and sidechain amino nitrogen of **3au**, and other between backbone carbonyl oxygen of GLY151 and sidechain amino-NH of **3au**. Anchoring due to these additional H-bonding interactions favoured hydrophobic interaction of 4-methoxy phenyl ring at 4^th^ position of thiazole ring with S2 (ASP75) subpocket residue along with NS2B (ASP81) residue shaping the S2 pocket in a closed active conformation. Interaction of both the phenyl rings thus kept the thiazole ring with hyradrazone linker showing hydrophobic interaction with S2 (ASN152) subpocket residues. S1 and S2 subpockets demands dibasic amino acid requirement that poses challenge in optimizing the PK profile of peptidic inhibitor design. Our compounds did not show any interaction with residues lining S1 subpocket (LEU115, SER163, and ILE165) except TYR150, basic requirement for S2 subpocket has been (partially) fulfilled by heterocyclic ring (thiazole) with hydrazone sidechain. This along with the fact that both the compounds displayed hydrophobic interaction with residues of catalytic triad (HIS51, ASP75, and SER135) justifies their inhibitory activity in-vitro. Fig. 3C and 3D displays comparative 2D-interaction plot for compounds **3aq** and **3au** with DENV NS2B-NS3 protease.

**Figure 3.**
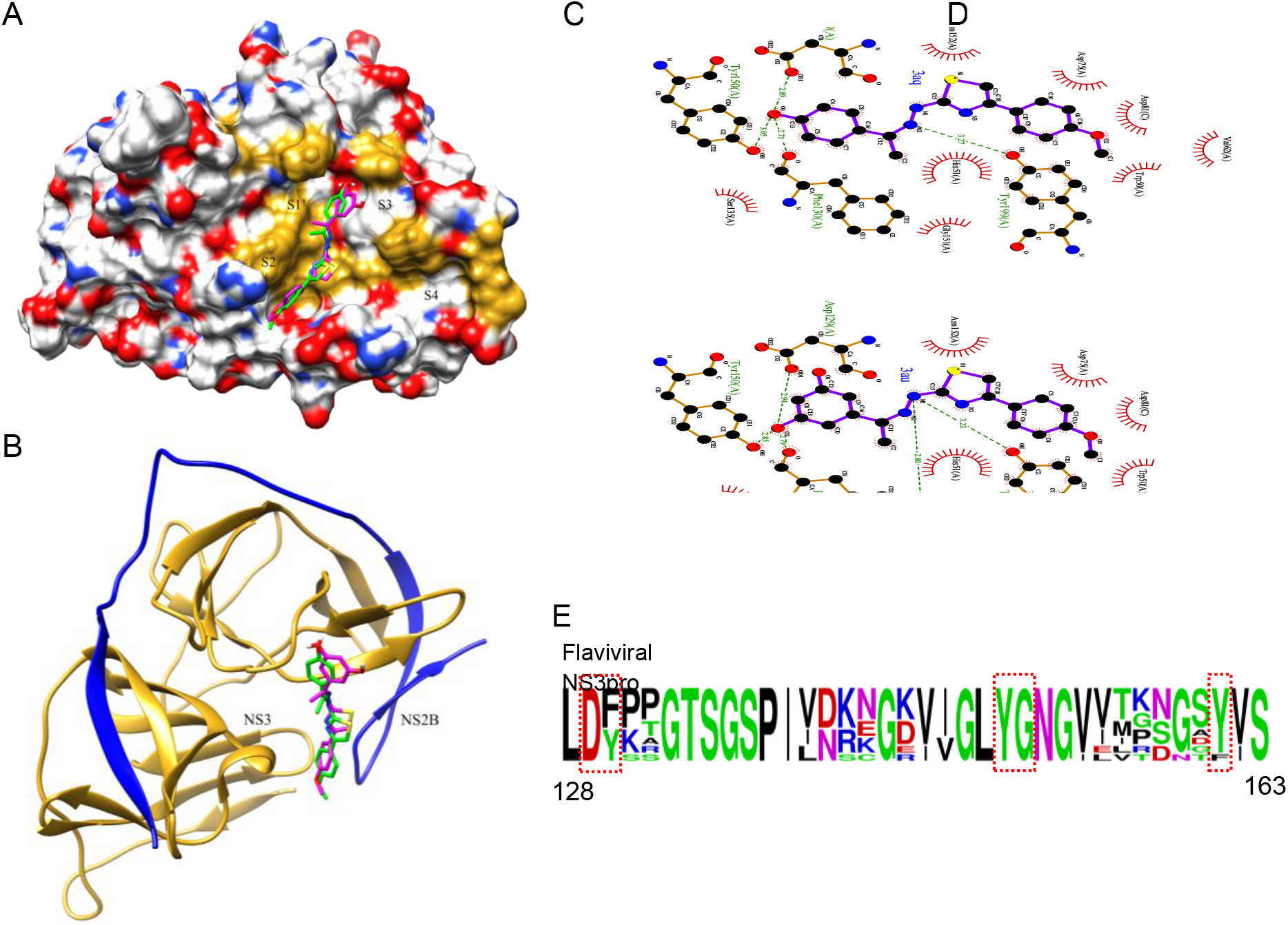
Docking image of compounds **3aq** (green) and **3au** (magenta) against DENV2 NS2B-NS3 protease crystal structure (PDB ID: 3U1I) (A) 3D representation of docking image in solvent-accessible surface colored by heteroatom while S1’,S2, S3, and S4 are the sub-pockets (residues lining the pockets colored golden). (B) The ribbon model depicts the protease accommodating ligands, aligned as shown in (A) NS3 colored golden and NS2B colored blue.. Images were generated using Chimera software. (C, D) Two-dimensional interaction diagrams illustrating the interactions of compounds **3aq** and **3au** with DENV2 NS2B-NS3 protease (PDB: 3U1I). (E) Logo plot showcasing amino acid frequencies in NS3 protease across various flaviviruses, emphasizing the conservation of crucial interacting residues: D129, F130, Y150, G151 and Y161.

## Detailed characterization of the inhibitory action of two potent compounds

As **3au** and **3aq** compounds emerged as the two most promising candidates, we characterized their NS2B-NS3 inhibitory activity in greater detail using the previously mentioned DENV NS2B-NS3 protease assay (Fig. 2D). Detailed kinetic experiments were performed to determine the kinetic parameters, inhibition constants, and inhibition mechanisms of these compounds. Various substrate concentrations were assessed with either the vehicle control or two different drug concentrations. Subsequently, enzymatic activity and corresponding reaction velocity were measured. Several inhibitory mechanisms, including competitive, uncompetitive, and mixed/non-competitive mechanisms, were explored for each drug. As shown in Tables A and B in Fig. 4, the presence of the compounds reduced both the K_m_ and V_max_ of the reaction, thereby indicating an uncompetitive mode of inhibition for both compounds. The double reciprocal plot (Lineweaver Burk plot) (Fig. 4C and 4D) also validates the mode of enzymatic inhibition against DENV NS2B-NS3 protease. This is also supported by the docking analysis, which showed no polar interaction with the triad site, similar to substrate backbone with partial occupation of S2 sub-pocket NS2B-NS3 protease (Fig. 3).

**Figure 4.**
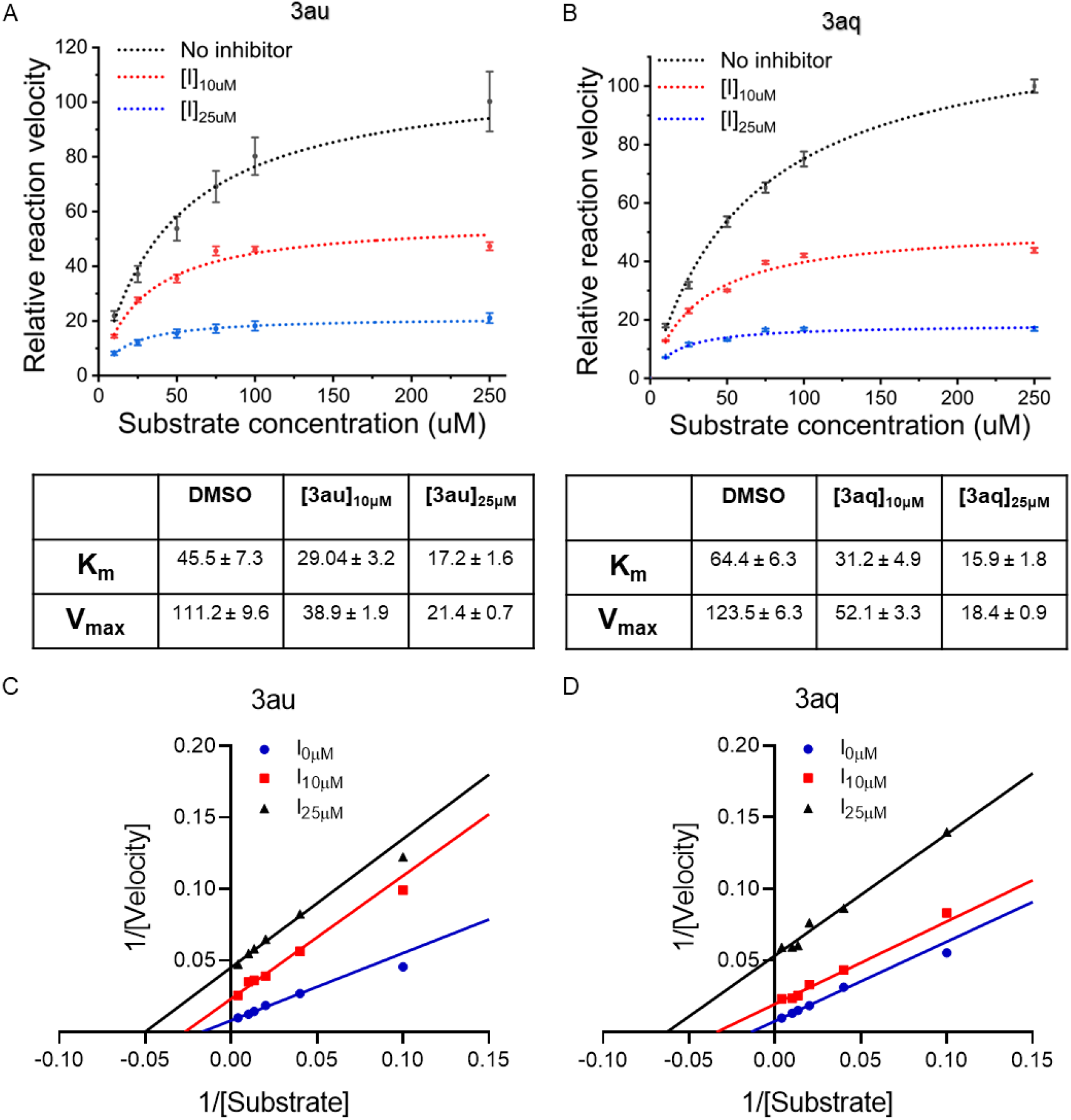
In-depth characterization of the inhibition pattern of top two potent compounds (A, B, C and D) Similar *in vitro* protease inhibition assay was performed in presence of two different concentrations of each compound using increasing substrate concentrations. NS2B-NS3 enzymatic kinetics was determined by fitting the nonlinear equation in Origin2023b software. Table representing two kinetics parameters, K_m_ and V_max_ of the control enzyme as well treated enzyme. The kinetic data were fitted into the Michaelis-Menten equation. (C and D) Furthermore, Lineweaver-Burk plot was done in Origin2023b.

## Anti-flaviviral activity of 3aq and 3au

*In vitro* studies suggested that **3au** and **3aq** can inhibit DENV NS2B-NS3 activity (Fig. 2–3, 4). Furthermore, our docking analysis revealed that these two compounds preferentially bind to several residues within and around the catalytic core of NS2B- NS3, most of which remain highly conserved across different flaviviruses (Fig. 3).(20) Hence, we aimed to assess the antiviral activity of these compounds. First, cytotoxic impact of these compounds was evaluated using MTT assay. Our data demonstrated that both compounds were non-toxic at concentrations of up to 10 μM. Although compound **3au** showed mild toxicity at 20 μM, **3aq** remained non-toxic at this concentration (Fig. 5A-5B). Subsequently, we evaluated the effectiveness of these compounds in inhibiting DENV2 and JEV replication by tracking viral genomic RNA replication in infected cells. BHK-21 cells were infected with the virus, either with or without the presence of these drugs, and harvested at various time points post-infection. The viral replication rate was assessed by quantifying the copy number of viral genomic RNA in the treated and control groups using quantitative real-time PCR (qRT-PCR). Our data suggested that both 3au and 3aq, at 10 μM concentrations, resulted in 90.1% and 95.9% reduction in viral RNA accumulation, respectively, during JEV infection. For DENV2 there was 47.1% and 67.6% reduction for 3au and 3aq respectively (Fig. 5C– 5D). Clearly, the compounds strong antiviral effects against JEV infection.

**Figure 5.**
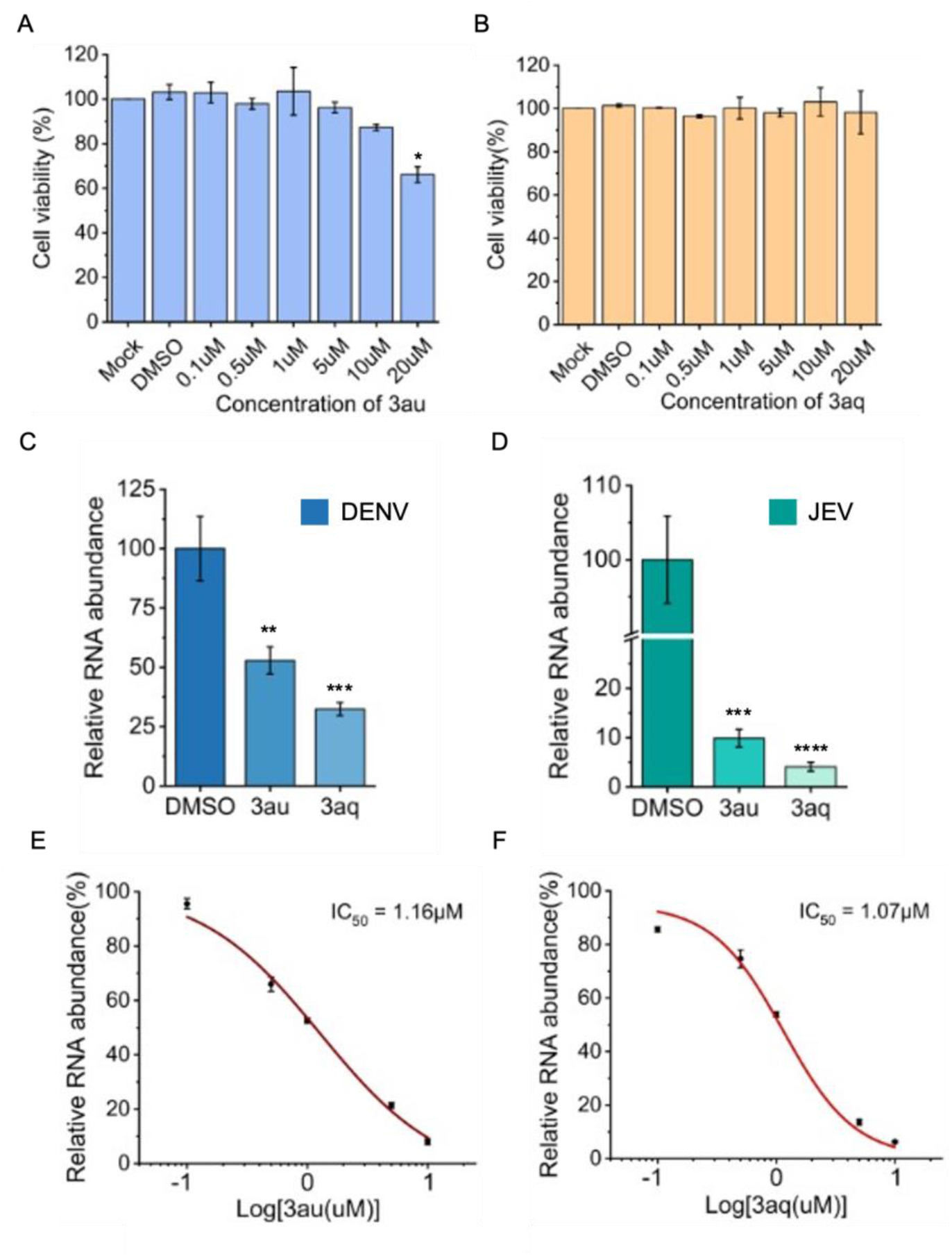
Anti-flaviviral activity of **3au** and **3aq** inhibiting DENV and JEV replication (A, B) BHK-21 cells were treated with different concentrations of the compounds (**3aq** and **3au**) for 24 h. MTT assay was performed to assess the cytotoxicity of the drugs. (C, D) BHK-21 cells were infected with DENV and JEV, respectively, in the absence or presence of 10 µM of each compound at a multiplicity of infection (MOI) of 0.1 and incubated for 24 hours at 37 °C. Reverse transcription and subsequent quantitative RT- PCR (qRT-PCR) were performed to measure the replication efficiency. Statistical significance was analysed by two-tailed equal variance Student’s t-test: *P<0.05, **P<0.01, ***P<0.001 and ****P<0.0001. (E & F) BHK-21 cells were infected with JEV virus inoculum with different concentrations of each compound at a multiplicity of infection (MOI) of 0.1 and incubated for 24 hours at 37 °C. Similar RNA quantification of the infected cells was conducted. Furthermore, the IC_50_ of the compounds against JEV virus was determined by fitting the nonlinear equation in Origin2023b software.

To this end, we investigated the antiviral activity of these compounds against JEV in detail. BHK-21 cells were infected with JEV (multiplicity of infection (MOI: 0.1) in the absence and presence of increasing concentrations of the compounds, followed by harvesting of the cells at 18 hpi. qRT-PCR was performed to monitor viral RNA accumulation as a proxy for viral replication. As shown in Fig. 5, both drugs showed a dose-dependent decrease in viral genomic RNA copy number, with more than 90% reduction observed with 10 μM concentration of each compound (Fig. 5E–5F). Further, as observed from the curve, the 50% inhibitory concentration of the compounds against JEV was determined to be 1.07 μM and 1.16 μM for **3aq** and **3au,** respectively (Fig. 5C–5D). These data confirmed the efficacy of the compounds in inhibiting JEV replication, possibly through direct interactions with NS2B-NS3 protein, thereby inhibiting its activity, as evidenced from the biochemical analyses. It is important to note that both compounds showed more prominent antiviral effect towards JEV in cell culture (IC50 1.16 μM for 3au and 1.07 μM for 3aq) than that could be predicted from the protease inhibitory activity evidenced from the biochemical analysis (IC50 20.76 μM for 3aq and 15.15 μM for 3aq). This indicated that apart from their role in blocking NS3- NS3B protease activity the compounds may impact additional viral machineries thereby blocks the overall viral propagation to an extent higher than expected.

## Compounds 3aq and 3au impact viral replication and accumulation of viral genomic RNA at late stages of infection

Flaviviral replication kinetics showed delayed accumulation of viral RNA at later time points during the infectious cycle(43). This is because the initial translation of the viral RNA genome leads to the production of a viral polyprotein, which, upon processing by the NS2B-NS3 protease, leads to the generation of viral RdRp (NS5) and other NS proteins necessary for RNA synthesis(13, 16). Thus, inhibition of the NS2B-NS3 protease by these two potential drugs should affect viral genomic RNA accumulation within infected cells only at later time points post-infection(43). To validate this hypothesis, we studied the effects of these drugs on JEV replication kinetics. Cells were infected with the virus in the absence or presence of **3aq** or **3au** (10 μM), harvested at different time points post-infection, and viral genomic RNA copies were estimated using qRT-PCR. Fig. 6A shows the relative levels of viral genomic RNA in treated versus control sets and Fig. 6B shows the absolute expression of viral RNA in the control and treated sets during the course of infection. As expected, the abundance of viral genomic RNA remained low at the early time points of infection (1 and 9 hpi) and subsequently showed a sharp increase at 18 hpi (Fig. 6B). Interestingly, both compounds showed a drastic effect on viral genomic RNA abundance only at the later time point post-infection (18 hpi), whereas no effect was observed at the early time points (1 and 9 hpi) (Fig. 6A– 6B). These data clearly demonstrate that **3aq** and **3au** exert their antiviral effects by blocking genomic RNA replication at later time points post-infection, which is an indirect effect of NS2B-NS3 protease inhibition, thereby blocking polyprotein processing and assembly of the viral RNA synthesis machinery. Together, our data not only established the antiviral activity the compounds against a DENV and JEV and also pointed out the mechanism of action of these drugs upon the specific steps of virus life cycle.

**Figure 6.**
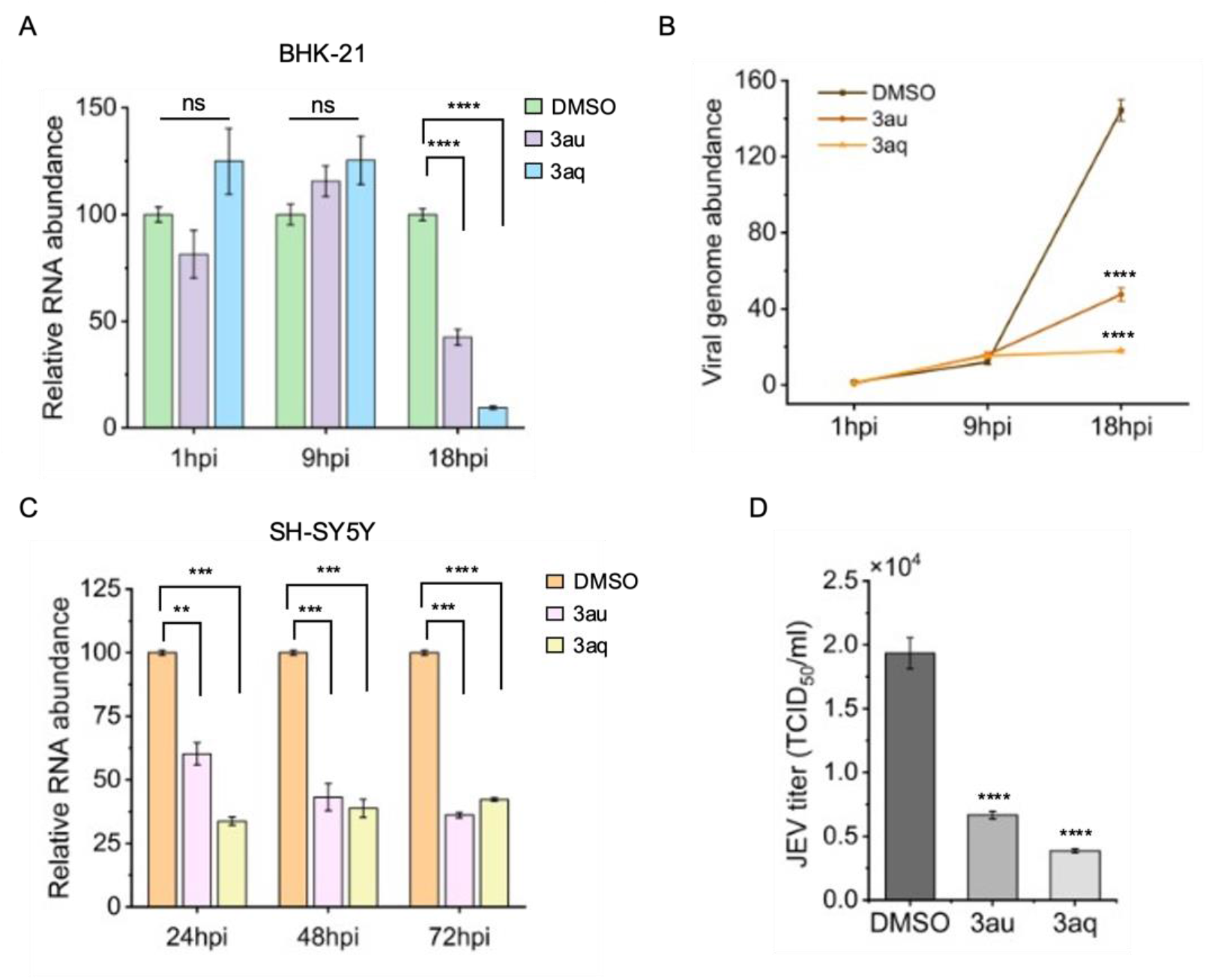
Detailed molecular mechanism of the inhibitory action of the compounds. (A) BHK-21 cells were infected with JEV virus (MOI=0.1) for 1, 9 & 18 h with or without 10 µM of compounds. (C) SH-SY5Y cells were infected with JEV virus (MOI=0.1) for 24, 48 & 72 h with or without 10 µM of compounds. (A & C) Total RNA extracted at indicated times, were evaluated by qRT-PCR. The data are expressed as fold changes of the viral RNA levels normalized to GAPDH control relative to the corresponding DMSO treated samples at each time point. (B) Total RNA abundance was also measured to understand the replication kinetics at different time points of infection. (D) BHK-21 cells were infected with JEV virus (MOI=0.1) for 18 h in presence of 10 µM of compounds and DMSO used as control set. The supernatant was collected to detect the JEVtiter using 50% tissue culture infective dose (TCID50) assays.Significance values were determined using Student’s t-Test. *P < 0.05, **P < 0.01, ***P < 0.001, and ****P < 0.0001.

To validate the antiviral effects of both compounds on JEV infectivity in a human cell line, we further studied the effect of the drugs on JEV replication kinetics in human neuroblastoma SH-SY5Y cells. Cells were infected with the virus in the absence or presence of **3aq** or **3au** (10 μM), and viral genomic RNA copies were quantified using qRT-PCR at 24, 48, and 72 hpi, as described previously. Considering that the compounds do not impact viral RNA accumulation at early time points (1 and 9 h.p.i) of post infection, these time points are excluded. qRT-PCR results demonstrated that **3au** and **3aq** treatment caused a sharp reduction in viral RNA levels at different time points, with 40% and 67% reduction for **3au** and **3aq**, respectively, at 24 hpi; 54% and 63% reduction for **3au** and **3aq**, respectively, at 48 hpi, and 64% and 58% reduction for **3au** and **3aq**, respectively, at 72 hpi (Fig. 6C). These results revealed that **3au** and **3aq** also exhibited antiviral activity against JEV production in SH-SY5Y cells, correlating with the antiviral effects shown in non-human cells (Fig. 5D). The antiviral activity of these two compounds was also determined by evaluating the extent of progeny virus production. Virus supernatants were harvested from JEV-infected BHK-21 cells, either treated with **3aq** or **3au** or with DMSO, at 18 hpi and virus titer was determined using TCID_50_ assay. Compound **3aq** caused the maximum reduction of virus titer from 1.93×10^4^ to 3.85×10^3^ TCID_50_/mL, whereas treatment of **3au** decreased the titer to 6.66×10^3^ TCID_50_/mL (Fig. 6D). Collectively, these infection data support the strong inhibition of JEV amplification by compound **3au** and **3aq.**

## Conclusion

With the increasing impact of flavivirus infections worldwide, a broad-spectrum antiviral agent targeting the highly conserved viral machinery is needed. The proteolytic processing of viral polyproteins is a key event that is imperative for the production of viral structural and non-structural proteins. This ensures the smooth occurrence of the subsequent steps of the viral life cycle, including genomic RNA replication, packaging into newly assembled viral capsids, and the final release of mature virion particles from the host cells (18, 37). Therefore, virus-encoded proteases have emerged as effective antiviral targets, not only for flaviviruses but also for other positive-sense RNA viruses, such as picornaviruses or coronaviruses(44). Interestingly, the catalytic core (catalytic triad) of the flavivirus NS3 protease (DENV, JEV, YFV, WNV, and ZIKV) shows high sequence and structural conservation, with similar sub-pocket residues and substrate amino acid preferences (at the P1 and P2 positions).(19, 20) Considering these factors, a previously reported retro-tripeptide (Weigel et al., 2015) was transformed into a thiazole heterocyclic template and derivatized to synthesize a series of thiazole compounds (**3a**-**3av**). These compounds were evaluated for their DENV NS2B-NS3 protease inhibitory activities using a high-throughput *in vitro* screening system(38–40). The top two compounds (**3aq** and **3au)** showing the highest inhibition in the *in vitro* screening were shortlisted for further characterization. Docking of these two compounds on the DENV NS2B-NS3 protease crystal structure (PDB ID: 3U1I) revealed that these compounds did interact with the catalytic triad (Fig. 2) and interacted with residues situated within sub-pockets S1’, S2 and S3. Interestingly, both compounds affected K_m_ and V_max_, implying uncompetitive inhibition of the NS2B-NS3 protease (Fig. 4).

Both compounds showed effective inhibition of DENV2 and JEV infection in BHK-21 cells, with low micromolar IC_50_ values (1.07 μM and 1.16 μM for compounds **3aq** and **3au**, respectively) specifically for JEV (Fig. 5). Interestingly, the residues targeted by both compounds remained conserved across a wide range of flaviviruses, including DENV, JEV, YFV, WNV, and ZIKV, indicating probability of antiviral activity against a wide range of flaviviruses. We also determined the inhibitory mechanisms of these compounds on viral replication. Time-course experiments showed that the rate of accumulation of viral genomic RNA by both these compounds remained unchanged at early time points (1 and 9 hpi), but was significantly reduced at later time points post-infection (18 hpi). These data confirm that the shortlisted compounds do not affect early events of the infectious cycle, such as receptor binding, endocytosis, uncoating, or release of viral genomic RNA within the host cellular cytoplasm. Antiviral activity correlates with NS2B-NS3 inhibitory activity, which affects polyprotein processing and eventually blocks viral RNA synthesis. It is interesting that the compounds showed more pronounce effect in inhibiting virus RNA synthesis in cell culture than the inhibitory effect showed in the context of in vitro protease activity assay. It is possible that along with the protease inhibitory function, these compounds may impact additional parts of the replication machinery and these two effects acts in synergism to show a strong antiviral effect against JEV.

SH-SY5Y cells are extensively used as an essential model for techniques focused on neurological illnesses during JEV or other flaviviral disease conditions. Taking advantage of SH-SY5Y cells as a neuronal-like *in vitro* model, numerous studies resulted in significant biological and pharmacological breakthroughs.(45–48) Study in human neuroblastoma SH-SY5Y cells also established the antiviral effects of these compounds against JEV (Fig. 6C). The TCID_50_ assay was used to measure the titer of JEV in the supernatant of infected BHK-21 cells at 18 hpi in the presence of both compounds. Again, both compounds showed promising effects by inhibiting viral amplification (Fig. 6D). Inhibition of viral titer correlated with the results of viral RNA synthesis upon treatment, entrenching the drastic effect of these compounds upon viral replication. Together, these data establish the newly synthesized **3aq** and **3au** compounds as inhibitors of DENV NS2B-NS3 protease, which has the potential to be developed as antivirals agents against multiple members of the Flaviviridae family.

## Experimental section

### General comments

All chemicals and reagents were purchased from commercial suppliers (CDH/Merck/Sigma-Aldrich/Alfa Aesar/Rankem) and were used without purification. TLC plates (Merck) were used to monitor the completion of the reactions in either iodine or UV chamber (at 254 nm). Melting points were determined using OPTIMELT (Stanford Research System, USA) and were uncorrected. All the final compounds and a few intermediates were characterized by ^1^H-NMR and ^13^C-NMR spectroscopy on a JEOL JNM-ECZ 400S/L1400 MHz NMR spectrometer with tetramethylsilane (TMS) as the internal standard and CDCl_3_ and DMSO-d_6_ were used as solvents. For the ^1^H-NMR spectra, the coupling constant (J) is expressed in hertz (Hz). The chemical shifts (δ) of NMR have been reported in parts per million (ppm) units relative to TMS. The splitting patterns are abbreviated as singlet (s), doublet (d), double doublet (dd), triplet (t), quartet (q), and multiplet (m). Mass spectra were recorded on a Thermo Scientific Ultimate 3000/Waters (Milford, USA) using electron spray ionization (ESI). The molecular docking studies were performed on MacBook Pro with Apple M2 chip and 8GB Memory running on macOS Sonoma 14.6.1. ChemDraw 19.1 (Perkin- Elmer Informatics) was used to draw ligand structures. The protein structure was downloaded from the Protein Data Bank (https://www.rcsb.org/).(49, 50) LigPlot + was used for visualization of the 2D ligand interactions, and LigPlot+ was used.(51)

### General procedure for the synthesis of (E)-2-(benzylidene/substituted benzylidene)-(1 (substituted-phenylethylidene)-hydrazine carbothioamide (2a-u)

To a solution of acetophenone/benzaldehyde (0.01 mol) and thiosemicarbazide (0.012 mol) in methanol (25 mL), a catalytic amount of acetic acid glacial (0.6-0.9 mL) was added. The mixture was stirred for a period–2-4 h at room temperature. After confirming the completion of the reaction by TLC, the solution was diluted with ice water (50 mL), and the obtained solid was filtered, washed with distilled water, dried, and recrystallized from hot methanol.

### General procedure for the synthesis of 2-[(2E)-2-benzylidene/-(1- phenylethylidene)-hydrazinyl]-4-phenyl-1,3-thiazole (3a-3av)

An appropriate amount of phenacyl bromide (0.01 mol) was then added to a solution of hydrazine carbothioamide (0.01 mol) in methanol (25 mL). The mixture was stirred for a period 30-90 min at room temperature. The completion of the reaction was confirmed by TLC and further the solution was diluted with ice water (50 mL), and the obtained solid was filtered, washed with distilled water, dried, and recrystallized from hot methanol.^30^

#### (E)-2-(2-(4-methylbenzylidene)hydrazinyl)-4-(3-nitrophenyl)-thiazole (**3a**)

Orange amorphous powder; yield (%): 70; mp (°C): 201-203; ^1^H-NMR (400 MHz,DMSO-D_6_): δ (ppm) 2.40 (s, 3H), 7.35 (s, 2H), 7.70 (d, J = 37.9 Hz, 4H), 8.40-8.10(m, 3H), 8.77 (s, 1H), 12.34 (s, 1H) ; ^13^C-NMR (101 MHz, DMSO-D_6_):δ (ppm) 21.7, 107.0, 120.6, 122.7, 127.0, 130.1, 130.9, 132.2, 136.9, 139.8, 142.5, 148.9,169.3; ESI-MS (m/z): 339.04 [M+H]^+^.

#### (E)-2-(2-(3,5-dimethylbenzylidene)hydrazinyl)-4-(3-nitrophenyl)thiazole (**3b**)

Pale yellow amorphous powder; yield (%): 98; mp (°C): 208-210; ^1^H-NMR (400 MHz,DMSO-D_6_): δ (ppm) 2.38 (s, 6H), 7.09 (s, 1H), 7.34 (s, 2H), 7.76 (d, J = 18.7 Hz,2H), 8.05-8.37 (m, 3H), 8.75 (s, 1H), 12.33 (s, 1H); ^13^C-NMR (101MHz, DMSO-D_6_): δ (ppm) 21.5, 107.0, 120.6, 122.7, 124.8, 130.9, 131.6, 132.2, 134.8,136.8, 138.5, 142.7, 148.7, 148.9, 169.2; ESI-MS (m/z): 353.05 [M+H]^+^.

#### (E)-2-(2-(2-methoxybenzylidene)hydrazinyl)-4-(3-nitrophenyl)-thiazole (**3c**)

Mustard yellow amorphous powder; yield (%): 59; mp (°C): 186-188; ^1^H-NMR (400MHz, DMSO-D_6_): δ (ppm) 3.85 (s, 3H), 7.08 (t, J = 7.4 Hz, 1H), 7.16 (d, J = 8.2 Hz,1H), 7.44 (t, J = 7.1 Hz, 1H), 7.71 (s, 1H), 7.77 (t, J = 7.7 Hz, 1H), 7.86 (d, J = 7.7Hz, 1H), 8.21 (d, J = 8.2 Hz, 1H), 8.34 (dd, J = 20.1, 8.0 Hz, 1H), 8.43 (s, 1H), 8.76 (d, J = 19.2 Hz, 1H), 12.32 (s, 1H); ^13^C-NMR (101 MHz, DMSO-D_6_):δ (ppm) 56.2, 106.8, 112.3, 120.4, 121.3, 122.5, 125.4, 130.7, 131.4, 132.1, 136.7, 137.7,148.6, 157.6, 169.1; ESI-MS (m/z): 355.11 [M+H]^+^.

#### (E)-2-(2-(2,5-dimethoxybenzylidene)hydrazinyl)-4-(3-nitrophenyl)thiazole (**3d**)

Orange amorphous powder; yield (%): 69; mp (°C): 173-175; ^1^H-NMR (400 MHz,DMSO-D_6_): δ (ppm) 3.81 (s, 6H), 6.96-7.05 (m, 2H), 7.30 (d, J= 2.7 Hz, 1H), 7.69 (d, J = 17.6 Hz, 2H), 8.15 (d, J = 7.7 Hz, 1H), 8.30 (d, J = 9.3Hz, 2H), 8.67 (s, 1H), 12.30 (s, 1H) ; ^13^C-NMR (101 MHz, DMSO-D_6_):δ (ppm) 55.9, 56.7, 107.0, 109.8, 113.8, 116.8, 120.5, 122.6, 123.4, 130.8, 132.2, 136.7,137.4, 148.7, 148.8, 152.1, 153.8, 169.1; ESI-MS (m/z): 385.18 [M+H]^+^.

#### (E)-2-(2-(3,4-dimethoxybenzylidene)hydrazinyl)-4-(3-nitrophenyl)thiazole (**3e**)

Orange amorphous powder; yield (%): 94; mp (°C): 174-176; ^1^H-NMR (400 MHz, DMSO-D_6_): δ (ppm) 3.77 (s, 6H), 6.98 (d, J = 8.2 Hz, 1H), 7.15 (d, J= 8.2 Hz, 1H), 7.24 (s, 1H), 7.60 (s, 1H), 7.68 (t, J = 8.0 Hz, 1H), 7.95 (s, 1H), 8.12(d, J = 8.2 Hz, 1H), 8.27 (d, J = 7.7 Hz, 1H), 8.64 (s, 1H), 12.14 (s, 1H) ; ^13^C-NMR (101 MHz, DMSO-D_6_): δ (ppm) 55.9, 56.1, 106.7, 108.8, 112.2,120.4, 120.9, 122.5, 127.6, 130.8, 132.1, 136.7, 142.4, 148.6, 148.8, 149.5, 150.7, 169.2; ESI-MS (m/z): 385.06 [M+H]^+^.

#### (E)-2-(2-(2-**fl**uorobenzylidene)hydrazinyl)-4-(3-nitrophenyl)thiazole (**3f**)

Lemon yellow amorphous powder; yield (%): 82; mp (°C): 188-190; 1H-NMR (400MHz, DMSO-D_6_): δ (ppm) 7.34 (s, 2H), 7.49 (s, 1H), 7.74 (t, J = 7.4 Hz, 2H), 7.92(s, 1H), 8.27 (t, J = 30.8 Hz, 3H), 8.72 (s, 1H), 12.52 (s, 1H); 13C-NMR(101 MHz, DMSO-D_6_): δ (ppm) 107.4, 116.5, 116.8, 120.6, 122.7, 125.6, 126.4, 130.9,131.7, 132.2, 134.8, 136.7, 148.9, 159.5, 162.0, 168.9; ESI-MS (m/z):342.98 [M+H]^+^.

#### (E)-2-(2-(3-chlorobenzylidene)hydrazinyl)-4-(3-nitrophenyl)thiazole (**3g**)

Pale yellow amorphous powder; yield (%): 96; mp (°C): 208-210; ^1^H-NMR (400 MHz,DMSO-D_6_): δ (ppm) 7.54 (s, 2H), 7.76 (d, J = 33.0 Hz, 4H), 8.12 (s, 1H), 8.24 (s,1H), 8.39 (s, 1H), 8.77 (s, 1H), 12.56 (s, 1H) ; ^13^C-NMR (101 MHz,DMSO-D_6_): δ (ppm) 107.3, 120.5, 122.6, 125.5, 126.0, 129.5, 130.8, 131.3, 132.1, 134.2,137.0, 140.4, 148.8, 168.9; ESI-MS (m/z): 359.11 [M+H]^+^.

#### (E)-2-(2-(2-bromobenzylidene)hydrazinyl)-4-(3-nitrophenyl)thiazole (**3h**)

Yellow amorphous powder; yield (%): 52; mp (°C):191-193; ^1^H-NMR (400 MHz,DMSO-D_6_): δ (ppm) 7.47 (t, J = 6.9 Hz, 1H), 7.61 (t, J = 7.1 Hz, 1H), 7.81-7.88 (m, 3H), 8.06 (d, J = 7.7 Hz, 1H), 8.30 (d, J = 7.7 Hz, 1H), 8.45 (d, J = 7.7 Hz, 1H), 8.52(s, 1H), 8.81 (s, 1H), 12.71 (s, 1H) ; ^13^C-NMR (101 MHz, DMSO-D_6_):δ (ppm) 107.6, 120.7, 122.8, 123.4, 127.3, 128.8, 131.0, 131.7, 132.3, 133.7, 133.9, 136.8,140.4, 149.0, 169.0; ESI-MS (m/z): 405.00 [M+2]^+^.

#### (E)-2-(2-(3-bromobenzylidene)hydrazinyl)-4-(3-nitrophenyl)thiazole (**3i**)

Yellow amorphous powder; yield (%): 89; mp (°C):195-197; ^1^H-NMR (400 MHz, DMSO-D_6_): δ (ppm) 7.48 (s, 1H), 7.66-7.94 (m, 5H), 8.11 (s, 1H), 8.32 (d, J = 64.3Hz, 2H), 8.77 (s, 1H), 12.53 (s, 1H); ^13^C-NMR (101 MHz, DMSO-D_6_):δ (ppm) 107.3, 120.5, 122.6, 125.9, 129.0, 130.8, 131.5, 132.1, 132.4, 136.6, 137.3, 140.3,148.8, 168.9; ESI-MS (m/z): 404.94 [M+2]^+^.

#### (E)-4-(4-chlorophenyl)-2-(2-(4-methylbenzylidene)hydrazinyl)thiazole (**3j**)

Off white amorphous powder; Yield (%): 86; mp (°C): 230-232; ^1^H-NMR (400 MHz,DMSO-D_6_): δ (ppm) 2.35 (s, 3H), 7.25-7.57 (m, 7H), 7.86-8.05 (m, 3H), 12.14 (s, 1H); ^13^C-NMR (101 MHz, DMSO-D_6_): δ (ppm) 21.7, 105.1, 127.0, 127.9,129.3, 130.1, 132.4, 132.6, 134.3, 139.7, 142.2, 150.0, 169.1; ESI-MS(m/z): 328.04 [M+H]^+^.

#### (E)-4-(4-chlorophenyl)-2-(2-(3,5-dimethylbenzylidene)hydrazinyl)thiazole (**3k**)

White amorphous powder; Yield (%): 61; mp (°C): 193-195; ^1^H-NMR (400 MHz,DMSO-D_6_): δ (ppm) 2.38 (d, J = 2.7 Hz, 6H), 7.10 (s, 1H), 7.34-7.56 (m, 5H), 8.00(d, J = 42.3 Hz, 3H), 12.22 (s, 1H); ^13^C-NMR (101 MHz, DMSO-D_6_):δ (ppm) 21.5, 105.1, 124.7, 127.9, 129.3, 131.6, 132.5, 134.2, 134.9, 138.5, 142.3, 169.0; ESI-MS (m/z): 342.04 [M]^+^.

#### (E)-4-(4-chlorophenyl)-2-(2-(2-methoxybenzylidene)hydrazinyl)thiazole (**3l**)

White amorphous powder; Yield (%): 66; mp (°C): 160-162; ^1^H-NMR (400 MHz,DMSO-D_6_): δ (ppm) 3.82 (s, 3H), 7.03 (dd, J = 24.2, 8.2 Hz, 2H), 7.35-7.44 (m, 4H),7.80 (dd, J = 32.7, 7.4 Hz, 3H), 8.33 (s, 1H), 12.12 (s, 1H) ; ^13^C-NMR(101 MHz, DMSO-D_6_): δ (ppm) 56.4, 105.1, 112.5, 121.5, 123.0, 125.6, 127.9, 129.3,131.5, 132.6, 134.3, 137.6, 150.0, 157.7, 169.1; ESI-MS (m/z): 344.11 [M+H]^+^.

#### (E)-4-(4-chlorophenyl)-2-(2-(2,5-dimethoxybenzylidene)hydrazinyl)thiazole(**3m**)

Dark brown crystalline powder; Yield (%): 52; mp (°C): 182-184; ^1^H-NMR (400 MHz,DMSO-D_6_): δ (ppm) 3.71 (s, 6H), 6.91-7.00 (m, 2H), 7.26-7.43 (m, 4H),7.83 (d, J = 8.2 Hz, 2H), 8.28 (s, 1H), 12.13 (s, 1H) ; ^13^C-NMR (101MHz, DMSO-D_6_): δ (ppm) 55.9, 56.7, 105.0, 109.7, 113.7, 116.6, 127.7, 129.1, 132.4,134.1, 137.1, 149.8, 152.1, 153.7, 168.9; ESI-MS (m/z): 374.12 [M+H]^+^.

#### (E)-4-(4-chlorophenyl)-2-(2-(3,4-dimethoxybenzylidene)hydrazinyl)thiazole (**3n**)

White amorphous powder; Yield (%): 96.21; mp (°C): 184-186; ^1^H-NMR (400 MHz,DMSO-D_6_): δ (ppm) 3.95 (s,6H), 7.15 (d, J = 8.2 Hz, 1H), 7.31 (d, J =7.7 Hz, 1H), 7.41 (s, 1H), 7.52 (s, 1H), 7.61 (d, J = 8.2 Hz, 2H), 8.01 (d, J = 8.2 Hz,2H), 8.11 (s, 1H), 12.22 (s, 1H) ; ^13^C-NMR (101 MHz, DMSO-D_6_): δ (ppm) 56.2, 104.9, 109.0, 112.3, 121.0, 127.9, 129.3, 132.6, 134.3, 142.3, 149.7, 150.0,150.8, 169.1; ESI-MS (m/z): 374.05 [M+H]^+^.

#### (E)-4-(4-chlorophenyl)-2-(2-(2-fluorobenzylidene)hydrazinyl)thiazole (**3o**)

Off white amorphous powder; Yield (%): 61; mp (°C): 176-178; ^1^H-NMR (400 MHz,DMSO-D_6_): δ (ppm) 7.27 (d, J = 7.7 Hz, 2H), 7.40-7.45 (m, 4H), 7.84 (d, J = 8.2Hz, 3H), 8.20 (t, J = 4.1 Hz, 1H), 12.34 (s, 1H); ^13^C-NMR (101 MHz,DMSO-D_6_): δ (ppm) 105.3, 116.4, 116.7, 122.4, 122.5, 125.5, 126.3, 127.8, 129.2,131.6, 131.6, 132.5, 134.0, 134.5, 149.9, 159.4, 161.9, 168.6; ESI-MS(m/z): 332.10 [M+H]^+^.

#### (E)-2-(2-(3-chlorobenzylidene)hydrazinyl)-4-(4-chlorophenyl)thiazole (**3p**)

White amorphous powder; Yield (%): 68; mp (°C): 189-191; ^1^H-NMR (400 MHz,DMSO-D_6_): δ (ppm) 7.39-7.45 (m, 5H), 7.59 (d, J = 7.1 Hz, 1H), 7.67 (s, 1H),7.85 (d, J = 8.8 Hz, 2H), 8.00 (s, 1H), 12.33 (s, 1H) ; ^13^C-NMR (101MHz, DMSO-D_6_): δ (ppm) 105.5, 125.7, 126.2, 127.9, 129.4, 129.6, 131.5, 132.7,134.2, 134.4, 137.3, 140.4, 150.0, 168.9; ESI-MS (m/z): 347.92 [M+H]^+^.

#### (E)-2-(2-(2-bromobenzylidene)hydrazinyl)-4-(4-chlorophenyl)thiazole (**3q**)

White amorphous powder; Yield (%): 81; mp (°C): 218-220; ^1^H-NMR (400 MHz,DMSO-D_6_): δ (ppm) 7.29-7.85 (m, 9H), 8.34 (s, 1H), 12.43 (s, 1H) ;^13^C-NMR (101 MHz, DMSO-D_6_): δ (ppm) 105.4, 123.1, 127.1, 127.8, 128.6, 129.2,131.4, 132.5, 133.7, 134.0, 140.0, 149.9, 168.5; ESI-MS (m/z): 394.06[M+2]^+^.

#### (E)-2-(2-(3-bromobenzylidene)hydrazinyl)-4-(4-chlorophenyl)thiazole (**3r**)

Cream white amorphous powder; Yield (%): 67; mp (°C): 199-201; ^1^H-NMR (400MHz, DMSO-D_6_): δ (ppm) 7.45-7.55 (m, 4H), 7.64 (d, J = 8.2 Hz, 1H), 7.73 (d, J= 7.7 Hz, 1H), 7.94 (t, J = 9.1 Hz, 3H), 8.08 (s, 1H), 12.42 (s, 1H);^13^C-NMR (101 MHz, DMSO-D_6_): δ (ppm) 105.4, 122.8, 125.9, 127.8, 128.9, 129.2,131.6, 132.3, 132.5, 134.0, 137.4, 140.1, 149.9, 168.7; ESI-MS (m/z):394.00 [M+2]^+^.

#### (E)-4-(3-nitrophenyl)-2-(2-(1-(o-tolyl)ethylidene)hydrazinyl)thiazole (**3s**)

Maroon amorphous powder; yield (%): 68; mp (°C): 158-160; ^1^H-NMR (400 MHz,DMSO-D_6_): δ (ppm) 2.38 (s, 3H), 2.53 (d, J = 36.3 Hz, 3H), 7.36 (d, J = 32.4 Hz,4H), 7.73 (d, J = 36.3 Hz, 2H), 8.29 (d, J = 65.4 Hz, 2H), 8.79 (s, 1H), 11.32 (s, 1H); ^13^C-NMR (101 MHz, DMSO-D_6_): δ (ppm) 18.7, 21.4, 107.0, 120.5,122.5, 126.3, 128.6, 128.7, 130.8, 131.4, 132.1, 134.7, 135.7,136.9, 139.5, 148.7, 150.1,170.7; ESI-MS (m/z): 353.11 [M+H]^+^.

#### (E)-4-(3-nitrophenyl)-2-(2-(1-(p-tolyl)ethylidene)hydrazinyl)thiazole (**3t**)

Brown crystalline powder; yield (%): 92; mp (°C): 215-217; ^1^H-NMR (400 MHz,DMSO-D_6_): δ (ppm) 2.31 (s, 3H), 3.18 (s, 3H), 7.21 (t, J = 5.6 Hz, 2H), 7.64-7.72 (m,4H), 8.21 (d, J = 67.6 Hz, 2H), 8.65 (s, 1H), 11.35 (s, 1H) ; ^13^C-NMR(101 MHz, DMSO-D_6_): δ (ppm) 14.6, 21.3, 107.3, 120.5, 122.5, 126.2, 129.6, 130.8,132.1, 135.6, 136.9, 138.9, 147.5, 148.8, 170.7; ESI-MS (m/z): 353.11[M+H]^+^.

#### (E)-4-(1-(2-(4-(3-nitrophenyl)thiazol-2-yl)hydrazono)ethyl)phenol (**3u**)

Reddish orange amorphous powder; yield (%): 82; mp (°C): 225-227; ^1^H-NMR (400MHz, DMSO-D_6_): δ (ppm) 2.32 (s, 3H), 6.85 (d, J = 8.8 Hz, 2H), 7.68 (d, J = 7.7 Hz,3H), 7.74 (t, J = 8.0 Hz, 1H), 8.17 (d, J = 7.7 Hz, 1H), 8.34 (d, J = 7.7 Hz, 1H), 8.75(s, 1H), 9.03 (s, 1H), 11.22 (s, 1H); ^13^C-NMR (101 MHz, DMSO-D_6_):14.6, 107.1, 115.7, 120.6, 122.4, 127.8, 129.2, 130.7, 132.1, 136.9, 147.9, 148.9, 158.9,170.1, 170.9; ESI-MS (m/z): 355.11 [M+H]^+^.

#### (E)-5-(1-(2-(4-(3-nitrophenyl)thiazol-2-yl)hydrazono)ethyl)benzene-1,3-diol (**3v**)

Orange yellow amorphous powder; yield (%): 87; mp (°C): 263-265; ^1^H-NMR (400MHz, DMSO-D_6_): δ (ppm) 2.37 (s, 3H), 6.35 (s, 1H), 6.77 (s, 2H), 7.80 (q, J = 8.2 Hz,2H), 8.33 (dd, J = 66.8, 7.4 Hz, 2H), 8.83 (s, 1H), 9.45 (s, 2H), 11.40 (s, 1H); ^13^C-NMR (101 MHz, DMSO-D_6_): 14.7, 103.7, 104.7, 107.3, 120.5, 122.5, 130.8,132.0, 136.9, 140.2, 147.5, 148.8, 158.8, 170.7; ESI-MS (m/z): 371.07[M+H]^+^.

#### (E)-2-(2-(1-(2-methoxyphenyl)ethylidene)hydrazinyl)-4-(3-nitrophenyl)thiazole (**3w**)

Pale yellow amorphous powder; yield (%): 95; mp (°C): 153-155; ^1^H-NMR (400 MHz,DMSO-D_6_): δ (ppm) 2.35 (s, 3H), 3.92 (s, 3H), 7.06-7.19 (m, 2H), 7.43 (d, J = 29.1Hz, 2H), 7.72-7.82 (m, 2H), 8.32 (dd, J = 66.8, 8.0 Hz, 2H), 8.81 (s, 1H), 11.34 (s,1H) ; ^13^C-NMR (101 MHz, DMSO-D_6_): 18.9, 56.2, 107.3, 112.3, 120.6,121.0, 122.6, 129.3, 129.9, 130.7, 130.9, 132.2, 148.9, 157.7, 170.8; ESI-MS(m/z): 369.12 [M+H]^+^.

#### (E)-2-(2-(1-(3-methoxyphenyl)ethylidene)hydrazinyl)-4-(3-nitrophenyl)thiazole (**3x**)

Yellow amorphous powder; yield (%): 71; mp (°C): 102-104; ^1^H-NMR (400 MHz,DMSO-D_6_): δ (ppm) 2.39 (s, 3H), 3.87 (s, 3H), 7.03 (s, 1H),7.41 (q, J = 2.9 Hz, 3H), 7.74-7.80 (m, 2H), 8.29 (d, J = 69.8 Hz, 2H), 8.79 (s, 1H),11.47 (s, 1H); ^13^C-NMR (101 MHz, DMSO-D_6_): δ (ppm) 14.7, 55.6,107.4, 111.7, 114.8, 118.8, 120.5, 122.5, 130.0, 130.8, 132.1, 136.9, 139.8, 147.0, 148.8,159.8, 170.7; ESI-MS (m/z): 369.18 [M+H]^+^.

#### (E)-2-(2-(1-(4-methoxyphenyl)ethylidene)hydrazinyl)-4-(3-nitrophenyl)thiazole (**3y**)

Orange amorphous powder; yield (%): 57; mp (°C): 164-166; ^1^H-NMR (400 MHz,DMSO-D_6_): δ (ppm) 2.29 (s, 3H), 3.78 (s, 3H), 6.97 (s, 2H), 7.62-7.74 (m, 4H), 8.22(d, J = 63.2 Hz, 2H), 8.71 (s, 1H), 11.25 (s, 1H); ^13^C-NMR (101MHz, DMSO-D_6_): δ (ppm) 14.6, 55.8, 107.2, 114.4, 121.1, 122.8, 127.8, 130.8, 132.1, 137.0, 147.4, 148.9, 160.5, 170.9; ESI-MS (m/z):369.11 [M+H]^+^.

#### (E)-2-(2-(1-(2,4-dimethoxyphenyl)ethylidene)hydrazinyl)-4-(3-nitrophenyl)thiazole (**3z**)

Yellow amorphous powder; yield (%): 94; mp (°C): 123-125; ^1^H-NMR (400 MHz, DMSO-D_6_) δ 2.33 (s, 3H), 3.89 (s, 6H), 6.68 (d, J = 23.6 Hz, 2H), 7.35 (s, 1H), 7.74 (d, J = 41.8 Hz, 2H), 8.30 (d, J = 65.4 Hz, 2H), 8.81 (s, 1H), 11.22 (s, 1H);^13^C-NMR (101 MHz, DMSO-D_6_): δ (ppm) 18.9, 56.3,99.4, 105.8, 107.1, 120.6, 122.1, 122.6, 130.7, 137.1, 148.8, 148.9, 149.8, 158.9, 161.8,170.9; ESI-MS (m/z): 399.13 [M+H]^+^.

#### (E)-2-(2-(1-(2,5-dimethoxyphenyl)ethylidene)hydrazinyl)-4-(3-nitrophenyl)thiazole (**3aa**)

Yellow amorphous powder; yield (%): 90; mp (°C): 170-172; ^1^H-NMR (400 MHz,DMSO-D_6_): δ (ppm) 2.41 (s, 3H), 3.90 (s, 6H), 7.11 (dd, J = 25.0, 15.1 Hz, 3H), 7.82(d, J = 32.4 Hz, 2H), 8.38 (d, J = 64.3 Hz, 2H), 8.88 (s, 1H), 11.42 (s, 1H); ^13^C-NMR (101 MHz, DMSO-D_6_): δ (ppm) 18.8, 56.1, 56.8, 107.4, 113.8,115.3, 115.6, 120.7, 122.7, 130.1, 131.0, 132.3, 137.1, 149.0, 149.5, 152.0, 153.6, 170.9; ESI-MS (m/z): 399.13 [M+H]^+^.

#### (E)-2-(2-(1-(3,4-dimethoxyphenyl)ethylidene)hydrazinyl)-4-(3-nitrophenyl)thiazole (**3ab**)

Yellow amorphous powder; yield (%): 76; mp (°C): 196-198; ^1^H-NMR (400 MHz, DMSO-D_6_) δ 2.37 (s, 3H), 3.86 (s, 6H), 7.03 (s, 1H), 7.35 (d, J = 8.2 Hz, 1H), 7.50 (s, 1H), 7.70 (s, 1H), 7.77 (t, J = 8.0 Hz, 1H), 8.28 (d, J = 68.7 Hz, 2H), 8.78 (s, 1H), 11.34 (s, 1H); ^13^C-NMR (101 MHz, DMSO-D_6_): δ (ppm) 14.5, 55.9, 56.1, 107.3,109.2, 111.8, 119.6, 120.6, 122.6, 130.8, 131.1, 132.2, 137.0, 147.4, 148.9, 149.1, 150.4,171.0; ESI-MS (m/z): 399.13 [M+H]^+^.

#### (E)-2-(2-(1-(4-chlorophenyl)ethylidene)hydrazinyl)-4-(3-nitrophenyl)thiazole (**3ac**)

Yellow amorphous powder; yield (%): 81; mp (°C): 177-179; ^1^H-NMR (400 MHz,DMSO-D_6_): δ (ppm) 2.39 (s, 3H), 7.54 (d, J = 8.8 Hz, 2H), 7.74-7.86 (m, 4H), 8.21(d, J = 8.2 Hz, 1H), 8.38 (d, J = 7.7 Hz, 1H), 8.78 (s, 1H), 11.52 (s, 1H); ^13^C-NMR (101 MHz, DMSO-D_6_): δ (ppm) 14.7, 107.5, 120.6, 122.6, 128.0,129.0, 130.8, 132.1, 134.0, 136.9, 137.2, 146.2, 148.9, 170.6; ESI-MS(m/z): 373.05 [M+H]^+^.

#### (E)-2-(2-(1-(4-fluorophenyl)ethylidene)hydrazinyl)-4-(3-nitrophenyl)thiazole (**3ad**)

Yellow amorphous powder; yield (%): 66; mp (°C): 202-204; ^1^H-NMR (400 MHz,DMSO-D_6_): δ (ppm) 2.43 (s, 3H), 7.36 (s, 2H), 7.85 (d, J = 56.1 Hz, 4H), 8.32(d, J = 67.6 Hz, 2H), 8.82 (s, 1H), 11.50 (s, 1H); ^13^C-NMR (101MHz, DMSO-D_6_): δ (ppm) 14.6, 107.4, 115.7, 116.0, 120.5, 122.5, 128.4, 130.7, 132.1,134.9, 136.9, 146.4, 148.8, 161.8, 164.3, 170.6; ESI-MS (m/z): 357.05 [M+H]^+^.

#### (E)-4-(4-chlorophenyl)-2-(2-(1-(p-tolyl)ethylidene)hydrazinyl)thiazole(**3ae**)

Light yellow amorphous powder; Yield (%): 91; mp (°C): 193-195; ^1^H-NMR (400MHz, DMSO-D_6_): δ (ppm) 2.29 (dd, J = 11.0, 7.1 Hz, 6H), 7.22 (d, J = 7.7 Hz, 2H),7.36-7.47 (m, 3H), 7.66 (t, J = 6.9 Hz, 2H), 7.87 (t, J = 7.7 Hz, 2H), 11.20 (s, 1H); ^13^C-NMR (101 MHz, DMSO-D_6_): δ (ppm) 14.7, 21.5, 105.5, 126.3,127.9, 129.3, 129.7, 132.5, 134.3, 135.8, 138.9, 147.3, 150.0, 170.7;ESI-MS (m/z): 342.17 [M+H]^+^.

#### (E)-4-(1-(2-(4-(4-chlorophenyl)thiazol-2-yl)hydrazono)ethyl)phenol (**3af**)

White amorphous powder; Yield (%): 66; mp (°C): 240-242; ^1^H-NMR (400 MHz,DMSO-D_6_): δ (ppm) 2.40 (s, 3H), 6.93 (s, 2H), 7.51 (s, 1H), 7.60 (d, J = 8.8 Hz, 2H),7.77 (d, J = 8.8 Hz, 2H), 8.03 (d, J = 8.2 Hz, 2H), 9.87 (s, 1H), 11.23 (s, 1H); ^13^C-NMR (101 MHz, DMSO-D_6_): δ (ppm) 14.7, 105.3, 115.9, 127.9, 129.3,129.5, 132.5, 134.5, 147.7, 150.1, 159.0, 170.9 Spectra B.125; ESI-MS (m/z): 344.11[M+H]^+^.

#### (E)-5-(1-(2-(4-(4-chlorophenyl)thiazol-2-yl)hydrazono)ethyl)benzene-1,3-diol (**3ag**)

White amorphous powder; Yield (%): 74; mp (°C): 241-243; ^1^H-NMR (400 MHz,DMSO-D_6_): δ (ppm) 2.31 (s, 3H), 6.33 (s, 1H), 6.75 (s, 2H), 7.47-7.56 (m, 4H), 7.97(d, J = 8.2 Hz, 2H), 9.44 (s, 2H), 11.26 (s, 1H) ; ^13^C-NMR (101 MHz, DMSO-D_6_): δ (ppm) 14.8, 103.8, 104.8, 105.5, 127.9, 129.3, 132.5, 134.3,140.4, 147.4, 150.0, 159.1, 170.7; ESI-MS (m/z): 360.07 [M+H]^+^.

#### (E)-4-(4-chlorophenyl)-2-(2-(1-(3-methoxyphenyl)ethylidene)hydrazinyl)thiazole (**3ah**)

Maroon amorphous powder; Yield (%): 64; mp (°C): 140-142; ^1^H-NMR (400 MHz,DMSO-D_6_): δ (ppm) 2.27 (s, 3H), 3.77 (s, 3H), 6.93 (s, 1H), 7.30-7.45 (m, 6H), 7.86(d, J = 8.8 Hz, 2H), 11.26 (s, 1H); ^13^C-NMR (101 MHz, DMSO-D_6_):δ (ppm) 14.6, 55.6, 105.5, 111.7, 114.7, 118.7, 127.8, 129.2, 130.0, 132.4, 134.2, 139.8,146.8, 150.0, 159.8, 170.5; ESI-MS (m/z): 358.11 [M+H]^+^.

#### (E)-4-(4-chlorophenyl)-2-(2-(1-(4-methoxyphenyl)ethylidene)hydrazinyl)thiazole (**3ai**)

Cream white amorphous powder; yield (%): 66; mp (°C): 201-203; ^1^H-NMR (400MHz, DMSO-D_6_): δ (ppm) 2.27 (s, 3H), 3.76 (s, 3H), 6.95 (d, J = 8.8 Hz, 2H), 7.35(s, 1H), 7.54-7.39 (2H), 7.71 (d, J = 8.8 Hz, 2H), 7.86 (d, J = 8.8 Hz, 2H), 11.15 (s,1H); ^13^C-NMR (101 MHz, DMSO-D_6_): δ (ppm) 14.6, 55.6, 105.3, 114.4, 127.7, 127.8, 129.2, 130.8, 132.4, 134.0, 147.5, 149.5, 160.9, 170.6; ESI-MS (m/z): 358.11 [M+H]^+^.

#### (E)-4-(4-chlorophenyl)-2-(2-(1-(2,4-dimethoxyphenyl)ethylidene)hydrazinyl)thiazole (**3aj**)

Light orange crystalline powder; Yield (%):79; mp (°C): 182-184; ^1^H-NMR (400 MHz,DMSO-D_6_): δ (ppm) 2.18 (s, 3H), 3.76 (s, 6H), 6.51-6.58 (m, 3H),7.21-7.43 (m, 3H), 7.84 (d, J = 8.8 Hz, 2H), 10.98 (s, 1H); ^13^C-NMR(101 MHz, DMSO-D_6_): δ (ppm) 18.7, 55.9, 99.2, 105.1, 105.6, 122.0, 127.8, 129.1,130.6, 132.4, 134.2, 149.6, 158.8, 161.6, 170.6; ESI-MS (m/z): 388.12[M+H]^+^.

#### (E)-4-(4-chlorophenyl)-2-(2-(1-(2,5-dimethoxyphenyl)ethylidene)hydrazinyl)thiazole (**3ak**)

Yellow amorphous powder; Yield (%): 46; mp (°C): 117-119; ^1^H-NMR (400 MHz,DMSO-D_6_): δ (ppm) 2.20 (s, 3H), 3.69 (s, 6H), 6.84-6.95 (m, 3H), 7.33-7.43 (m, 3H), 7.85 (d, J=11.0 Hz, 2H), 11.11 (s, 1H); ^13^C-NMR (101 MHz, DMSO-D_6_): δ (ppm) 18.6, 56.0, 56.6, 105.3, 113.5, 115.0, 115.4, 127.8,129.2, 130.0, 132.4, 134.2, 149.0, 150.1, 151.8, 153.4, 153.9, 169.5, 170.5; ESI-MS (m/z): 388.18 [M+H]^+^.

#### (E)-4-(4-chlorophenyl)-2-(2-(1-(3,4-dimethoxyphenyl)ethylidene)hydrazinyl)thiazole (**3al**)

Cream white amorphous powder; Yield (%): 80; mp (°C): 166-168; ^1^H-NMR (400MHz, DMSO-D_6_): δ (ppm) 2.43 (s, 3H), 3.94 (d, J = 11.0Hz, 6H), 7.12 (d, J = 8.2Hz, 1H), 7.41-7.62 (m, 5H), 8.03 (d, J = 8.8 Hz, 2H), 11.32 (s, 1H);^13^C-NMR (101 MHz, DMSO-D_6_): δ (ppm) 14.5, 56.2, 105.4, 109.3, 111.8, 119.5,127.9, 129.3, 131.2, 132.5, 134.4, 147.2, 149.2, 150.4, 170.8; ESI-MS(m/z): 388.18 [M+H]^+^.

#### (E)-4-(4-chlorophenyl)-2-(2-(1-(4-chlorophenyl)ethylidene)hydrazinyl)thiazole (**3am**)

White amorphous powder; Yield (%): 59; mp (°C): 194-196; ^1^H-NMR (400 MHz,DMSO-D_6_): δ (ppm) 2.65 (s, 3H), 7.61-8.05 (m, 9H), 11.52 (s, 1H);^13^C-NMR (101 MHz, DMSO-D_6_): δ (ppm) 14.5, 105.6, 127.8, 127.9, 129.0, 129.2,132.4, 133.9, 134.2, 137.2, 145.9, 150.0, 170.3; ESI-MS (m/z): 361.99[M]^+^.

#### (E)-4-(4-chlorophenyl)-2-(2-(1-(4-fluorophenyl)ethylidene)hydrazinyl)thiazole (**3an**)

White amorphous powder; Yield (%): 80; mp (°C): 200-202; ^1^H-NMR (400 MHz,DMSO- D_6_): δ (ppm) 2.35 (s, 3H), 7.29 (t, J = 8.8 Hz, 2H), 7.43-7.51 (m, 3H),7.84-7.94 (m, 4H), 11.33 (s, 1H) ; ^13^C-NMR (101 MHz, DMSO-D_6_):δ (ppm) 14.8, 105.6, 116.1, 127.9, 128.5, 129.3, 132.6, 134.4, 135.1, 146.3, 150.1, 162.0,164.4, 170.6; ESI-MS (m/z): 346.04 [M+H]^+^.

#### (E)-4-(1-(2-(4-phenylthiazol-2-yl)hydrazono)ethyl)phenol (**3ao**)

Cream white amorphous powder; yield (%): 55; mp (°C): 243-245 [Lit.182 257-269];^1^H- NMR (400 MHz, DMSO-D_6_): δ (ppm) 2.24 (s, 3H), 6.76-7.83 (m, 11H), 9.68 (s,1H), 11.01 (s, 1H); ^13^C-NMR (101 MHz, DMSO-D_6_): δ (ppm) 14.5,104.3, 115.7, 126.0, 127.9, 129.1, 135.4, 147.3, 151.2, 158.8, 170.5;ESI-MS (m/z): 310.04 [M+H]^+^.

#### (E)-4-(1-(2- (4-(p-tolyl)thiazol-2-yl)hydrazono)ethyl)phenol **(3ap)**

Cream colored amorphous powder; yield (%): 51; mp (°C): 205-207; ^1^H-NMR (400MHz, DMSO-D_6_): δ (ppm) 2.25 (d, J = 22.5 Hz, 6H), 6.76 (d, J = 8.2 Hz, 2H), 7.16(s, 3H), 7.66 (dd, J = 53.6, 8.0 Hz, 4H), 9.67 (s, 1H), 10.98 (s, 1H);^13^C-NMR (101 MHz, DMSO-D_6_): δ (ppm) 14.4, 21.3, 103.3, 115.7, 126.0, 127.7,129.4, 129.7, 132.8, 137.2, 147.3, 158.8, 170.4; ESI-MS (m/z): 324.01[M+H]^+^.

#### (E)-4-(1-(2-(4-(4-methoxyphenyl)thiazol-2-yl)hydrazono)ethyl)phenol (**3aq**)

Dark brown crystalline powder; yield (%): 38; mp (°C): 170-172; ^1^H-NMR (400 MHz,DMSO-D_6_): δ (ppm) 2.22 (s, 3H), 3.74 (s, 3H), 6.76 (d, J = 8.8 Hz, 2H), 6.93 (d, J =8.8 Hz, 2H), 7.07 (s, 1H), 7.59 (d, J = 6.6 Hz, 2H), 7.76 (d, J = 8.8 Hz, 2H), 9.67 (s,1H), 10.96 (s, 1H); ^13^C-NMR (101 MHz, DMSO-D_6_): δ (ppm) 14.4,55.6, 102.1, 114.5, 115.7, 127.3, 127.7, 129.4, 131.2, 147.3, 158.8, 159.2, 170.4; ESI-MS (m/z): 340.01 [M+H]^+^.

#### (E)-4-(1-(2-(4-(4-fluorophenyl)thiazol-2-yl)hydrazono)ethyl)phenol (**3ar**)

Buff colored amorphous powder; yield (%): 39; mp (°C): 235-237; ^1^H-NMR (400 MHz,DMSO-D_6_): δ (ppm) 2.23 (s, 3H), 6.76 (d, J = 8.8 Hz, 2H), 7.21 (q, J = 8.6 Hz, 3H),7.59 (d, J = 8.8 Hz, 2H), 7.86 (d, J = 5.5 Hz, 2H), 9.68 (s, 1H), 11.02 (s, 1H); ^13^C-NMR (101 MHz, DMSO-D_6_): δ (ppm) 14.5, 104.0, 115.7, 127.7, 129.4,132.1, 147.4, 150.1, 158.8, 160.9, 163.3, 170.6; ESI-MS (m/z): 328.00[M+H]^+^.

#### (E)-5-(1-(2-(4-phenylthiazol-2-yl)hydrazono)ethyl)benzene-1,3-diol (**3as**)

Orange yellow amorphous powder; yield (%): 37; mp (°C): 238-240; ^1^H-NMR (400MHz, DMSO-D_6_): δ (ppm) 2.19 (s, 3H), 6.20 (s, 1H), 6.62 (s, 2H), 7.24-7.37 (m, 4H),7.84 (d, J = 7.7 Hz, 2H), 9.29 (s, 2H), 11.11 (s, 1H); ^13^C-NMR (101MHz, DMSO-D_6_): δ (ppm) 14.7, 103.6, 104.6, 126.0, 128.0, 129.2, 135.4, 140.3, 147.0,151.2, 158.8, 170.4; ESI-MS (m/z): 326.04 [M+H]^+^.

#### (E)-5-(1-(2-(4-(p-tolyl)thiazol-2-yl)hydrazono)ethyl)benzene-1,3-diol (**3at**)

White amorphous powder; yield (%): 47; mp (°C): 135-137; ^1^H-NMR (400 MHz,DMSO- D_6_): δ (ppm) 2.23 (d, J = 39.0 Hz, 6H), 6.20 (s, 1H), 6.62 (s, 2H), 7.18 (d, J= 6.6 Hz, 3H), 7.72 (s, 2H), 9.29 (s, 2H), 11.08 (s, 1H); ^13^C-NMR (101MHz, DMSO-D_6_): δ (ppm) 14.7, 21.3, 103.6, 104.6, 126.0, 129.7, 132.7, 137.2, 140.3,146.9, 151.2, 158.8, 170.3; ESI-MS (m/z): 340.08 [M+H]^+^.

#### (E)-5-(1-(2-(4-(4-methoxyphenyl)thiazol-2-yl)hydrazono)ethyl)benzene-1,3-diol (**3au**)

Brown amorphous powder; yield (%): 51; mp (°C): 244-246; ^1^H-NMR (400 MHz,DMSO- D_6_): δ (ppm) 2.18 (s, 3H), 3.74 (s, 3H), 6.20 (s, 1H), 6.62 (s, 2H), 6.93 (d, J =8.8 Hz, 2H), 7.10 (s, 1H), 7.76 (d, J = 8.8 Hz, 2H), 9.29 (s, 2H), 11.07 (s, 1H); ^13^C-NMR (101 MHz, DMSO-D_6_): δ (ppm) 14.7, 55.6, 102.4, 103.6, 104.6, 114.5, 127.4,128.2, 140.3, 147.0, 150.8, 158.8, 159.3, 170.3; ESI-MS (m/z): 356.15[M+H]^+^.

#### (E)-5-(1-(2-(4-(4-fluorophenyl)thiazol-2-yl)hydrazono)ethyl)benzene-1,3-diol (**3av**)

Light yellow amorphous powder; yield (%): 33; mp (°C): 140-142; ^1^H-NMR (400 MHz,DMSO-D_6_): δ (ppm) 2.18 (s, 3H), 6.20 (d, J = 2.2 Hz, 1H), 6.61 (s, 2H), 7.18-7.27 (m,3H), 7.86 (d, J = 5.5 Hz, 2H), 9.15 (s, 2H), 11.11 (s, 1H); ^13^C-NMR(101 MHz, DMSO-D_6_): δ (ppm) 14.7, 103.7, 104.7, 115.9, 128.0, 131.9, 140.2, 147.2,158.8, 160.9, 163.3, 170.5; ESI-MS (m/z): 344.00 [M+H]^+^.

## Cells and viruses

Baby Hamster Kidney Fibroblast cells (BHK-21 cell line) were maintained in Dulbecco’s modified Eagle’s medium (DMEM) supplemented with 10% fetal bovine serum (FBS) and penicillin and streptomycin antibiotics (Gibco) at 37 °C in 5% CO_2_. *JEV* strain Vellore P20778, was used to infect BHK-21 cells.

## DENV2 NS2B-NS3 protease expression and purification

The pET28a plasmid encoding the NS2B/NS3 protease sequence from DENV2 was transformed into *Escherichia coli* BL21(DE3) cells. DENV2 NS2B-NS3 protease expression plasmids were grown in Luria-Bertani (LB) medium containing 50 μg/mL kanamycin at 37 °C until OD_600_ reached 0.6. First, the cultures were incubated for 2-3 h at 37 °C until an OD_600_ of 0.6–0.8 was reached. isopropyl β-d-1-thiogalactopyranoside (IPTG) was added at a final concentration of 1 mM. The cells were incubated for 5 h at 30 °C with shaking at 100 rpm to induce protein expression.

The cells were harvested by centrifugation at 4500 rpm (REMI NEYA16 centrifuge) for 15 min at 4 °C, and the pellets were frozen in liquid nitrogen and stored at -80 °C. The cell pellets were thawed and completely resuspended in lysis buffer (25 mM Tris pH 7.9, 0.5 M NaCl, 5 mM imidazole, 5% glycerol). After resuspension, the solution was maintained on ice. For purification, the cells were lysed by sonication using an Ultrasonic Cell Disruptor. The lysate was centrifuged at 15,000×g for 40 min at 4 °C. Prior to Ni^2+^ affinity chromatography, DNAse (Sigma-Aldrich) was added to the supernatant. The soluble 6x-His-NS2B-NS3 protease in its native form was filtered, batch-bound to 2 mL Ni^2+^-NTA (Qiagen) resin (pre-equilibrated with column buffer), and incubated for 1 h at 4 °C. The resin was removed from the unbound fraction by centrifugation and the resin containing the bound protein was collected and loaded into columns (Bio-Rad). The packed column was washed extensively with 10 mL wash buffer containing 5, 10, and 50 mM imidazole. The recombinant protein was eluted using elution buffer (lysis buffer containing 250 mM imidazole). The purified protein was analyzed using 12% SDS-PAGE. The fractions with the highest amount of pure protein were desalted and concentrated with a centrifugal concentrator (Sartorius: 10 kDa MWCO) and then washed 4-5 times with 40 mL of storage buffer (100 mM Tris, pH 7.9, 50 mM NaCl, and 50% glycerol) at 4 °C to remove all imidazole. The protein was distributed into 50 μL aliquots and frozen in liquid nitrogen for storage at -80 °C for further use in dengue protease activity and inhibition studies.

## NS2B-NS3 protease steady-state kinetics

A similar approach was used to analyze the enzymatic kinetics of protease inhibition. The reaction progress was monitored by the release of free aminomethylcoumarin (AMC), as previously described. The substrate was incubated with or without the inhibitor in the aforementioned reaction buffer, in the presence of varying concentrations of the substrate. The reactions were initiated by the addition of the NS2B-NS3 protease. Protease activity was monitored for 1 h based on the initial rate of increase in fluorescence intensity at 460 nm, with an excitation wavelength of 380 nm. The results were analyzed using Origin Pro 2023b and the Michaelis-Menten equation to obtain the apparent Michaelis-Menten constants and maximal velocities. All assays were performed in triplicates. For each tested compound, two different concentrations of the inhibitor (0, 10, and 25 μM) were used at various concentrations (0-250 μM). The mechanism of inhibition was determined by observing the deviations in the apparent K_m_ and V_max_ values.

## NS2B-NS3 Protease assay

The selected compounds were analyzed using *in vitro* protease assays performed in black 96-well plates. Standard reaction mixtures (60 μL) containing 50 mM Tris·HCl (pH 8.5), 1 mM CHAPS, 20% glycerol, 10 nM DENV2 NS2B-NS3 protease, 10 μM inhibitor (dissolved in DMSO) or DMSO and 10.0 μM tetra-peptide substrate trifluoroacetate-Nle-Lys-Arg-Arg-AMC (ALFA Chemicals) were added to the reaction mixture and incubated for 1 h at room temperature. All assays were performed in triplicates. The release of free AMC was measured using a Varioskan™ LUX multimode microplate reader (Thermo Fisher Scientific) at excitation and emission wavelengths of 380 nm and 460 nm, respectively. Control fluorescence values obtained in the absence of inhibitors were considered 100%, and those obtained in the presence of inhibitors were calculated as the percentage of inhibition of the control using Microsoft Excel and plotted using Origin Pro 2023b. The background AMC in the absence of protease was subtracted before data analysis. IC_50_ values were calculated using Origin Pro 2023b.

## MTT assay

The MTT assay was performed to determine cellular metabolic activity as an indicator of cell viability, proliferation, and cytotoxicity. Thus, to determine whether these compounds had any toxic effects on cellular viability, an MTT assay was performed following the protocol described previously by Mosmann (1983)(52). Briefly, BHK-21 cells were seeded into 96-well plates at a density of 20,000 cells/well. Almost 16-18 h later, the cells were treated with different concentrations of the compounds (**3au** and **3aq**) for 24 h at 37 °C in 5% CO_2_. Thereafter, the medium was aspirated and 100 μL of MTT reagent (5 mg/mL, SRL) in PBS was added to the cells and incubated for 3 h at 37 °C. Formazan crystals were formed in each well and resuspended in 100 μL of dimethyl sulfoxide (DMSO) (Sigma, USA) at room temperature for 20 min to completely dissolve the crystals. The absorbance of the solution in a 96-well plate was measured at 595 nm using a microplate reader (Epoch 2; BioTek Instruments). Each experiment was performed in triplicate and the percentage of cell viability was determined as the ratio of the absorbance values of the experimental samples to the control solvent (DMSO) on the same plate.

## JEV and DENV infection and compound treatment

BHK-21 cells were seeded in a 24-well cell culture plate at a density of (0.25 × 10^6^ cells/well) with regular growth medium containing DMEM + 10% FBS one day prior to infection in triplicate. Once the cells reached 85-90% confluence, they were infected with JEV virus inoculum comprising DMEM + 2% FBS and 10 µM of the compound at an MOI of 0.1. Cells were incubated for 1.5 h at 37 °C with the virus-compound inoculum, with gentle rocking every 10 min to ensure even coverage and prevent the cellular monolayer from drying. Finally, the inoculum was replenished with fresh culture media comprising DMEM + 2% FBS and the compound, and then kept in an incubator in a humidified chamber at 37 °C until harvest (18 h). The cell culture supernatant was aspirated and washed with PBS, and the cellular monolayer was used for viral RNA isolation.

## RT-PCR and quantitative PCR

Total cellular RNA was extracted from BHK-21 cells using TRIzol reagent, according to the manufacturer’s instructions (Ambion Life Technologies, USA). RNA quantity and integrity were determined prior to use on a NanoDrop spectrophotometer (Thermo Fisher Scientific). 1 μg of total RNA from each experimental sample was used to synthesize cDNA by reverse transcriptase (RT) by incubating the RNAs with Moloney murine leukemia virus (M-MuLV) RT, with specific RT primer, 5’- ACAGTCTTTCCTTCTGCTGCAGGTCT-3’ (T_m_ = 63 °C), and deoxynucleoside triphosphates (dNTPs) (New England Biolabs) at 42 °C for 50 min, followed by inactivation at 70 °C for 15 min.

To quantify the viral RNA copy number, quantitative RT-PCR (qRT-PCR) was performed in duplicate using iTaq Universal SYBR Green Supermix (Bio-Rad). The cDNAs used for qPCR analysis was prepared using Power SYBR Green PCR Master Mix (Bio-Rad, # 172-5125) in a QuantStudio5 Applied Biosystems real-time PCR system. For qPCR amplification of the JEV amplicon size of 202 bp, the sequences of the forward and reverse primer pairs used were 5’- GAGAGAGGCGCGCCAGTGGAAGGCTCAGGCGTCCAAAAG-3’ (T_m_ = 65 ⁰C) and 5’- ACAGTCTTTCCTTCTGCTGCAGGTCT-3.’ (T_m_ = 63 ⁰C) and for DENV amplicon of 163 bp, the sequences of the forward and reverse primer pair used were 5’- TAGCGGTTAGAGGAGACCCCTC-3’ (T_m_ = 61 ⁰C) and 5’-GAGACAGCAGGATCTCTGGTCT-3’ (T_m_ = 59 ⁰C) The viral amplicon levels were normalized to those of mouse/human glyceraldehyde 3-phosphate dehydrogenase (GAPDH) using the forward and reverse primers 5’- CATCACTGCCACCCAGAAGACTG-3’ and 5’-ATGCCAGTGAGCTTCCCGTTCAG-3’.

The average threshold cycle (*Ct*) values were converted to viral copy numbers by comparison with a standard plot established using known amounts of JEV cDNA. ΔCt = (*Ct* target mean *Ct* of reference genes). Delta *Ct* for both control and compound treated samples was determined and subsequently analyzed by determining the delta *Ct* to determine the mRNA expression of target genes in control vs. treated samples.

## Determination of virus titer by TCID_50_ assay

During JEV infection, the viral supernatant was collected and used in the TCID_50_ assay to determine the virus titer. 3×10^4^ BHK-21 cells per well) were seeded in a 96-well cell culture microtiter plate the day before titration. The virus stock was diluted with infection media (DMEM + 2% FBS) to attain virus dilutions of 1×10^-2^ to 6.4×10^-7^ in 12 parallels with the last column of the 96-well plate serving as virus-free cell control. On the day of the assay, when the cell had reached 90-95% confluency, the growth medium was aspirated using a multichannel pipette, and 100 µL of virus inoculum of different dilutions was transferred to each well, keeping the last well only in infection media without virus. The cells were then incubated at 37 °C in a humidified 5% CO2 chamber. The 96-well plates were examined under a light microscope 48 hpi for signs of CPE. The infection medium was then removed, and the cells were rinsed with PBS to remove dead cells from the well surface. Cells were fixed with 70% ethanol for 20 min and counterstained with 2% crystal violet, followed by scoring of each well as positive or negative for viral growth. TCID_50_ was calculated using the Spearman and Karber methods.

## Molecular modeling

The crystal structure of DENV3 protease (PDB ID: 3U1I)(42) was downloaded from the Protein Data Bank (PDB). Chain C (NS2B) and chain D (NS3) were extracted, three residues (V36I, T115L, and K157R) were mutated, and renumbered to mimic DENV2 NS3 protease. Further it was prepared using Dockprep utility in UCSF Chimera-1.18(53, 54) and saved as protein.mol2. Protein preparation includes a series of steps, such as water molecule deletion, addition of hydrogens, assignment of atom types (AD4), Gasteiger charge addition, application of selected flips to residues, and analysis of all-atom contacts. ChemDraw-19.1 was used to draw the 3D-geometry optimization and energy minimization of the most potent molecules (**3aq** and **3au**). Energy-minimized structures were saved in ligand.pdb format and further prepared using Dockprep utility in UCSF Chimera-1.18 and saved as ligand.mol2. Autodock Vina(55, 56) utility in UCSF Chimera-1.18 was used to perform docking. Grid centre (35.96, -41.97, 46.10) was specified using the cocrystallized ligand (OAR) with box size of 20 x 20 x 20, energy range of 3, exhaustiveness of 8 and number of modes 10. Docked conformers were then analyzed using View dock utility in UCSF Chimera-1.18. Complex files were saved as complex.pdb for generating 2D-interaction plots using Ligplot+(51).

## Acknowledgement

SM, DS acknowledges Science and Engineering Research Board, Govt. of India for Junior Research Fellowship (JRF) under Core Research Grant (EMR/2016/005711) sanctioned to BNS, VJ, and BS. IDJ acknowledges Science and Engineering Research Board (SERB) for National Post Doctoral Fellowship (<u>PDF/2023/000284</u>) awarded to her. A.M. acknowledges financial support from Science and Engineering Research Board (SERB-DST: CRG/2022/003628).

